# Analytical model of the selective cutting mechanism used by ovipositors of sawflies

**DOI:** 10.1101/2025.06.15.659745

**Authors:** Martí Verdaguer Mallorquí, Julian F.V. Vincent, Andrew Liston, Vladimir Blagoderov, Marc P.Y. Desmulliez

## Abstract

Sawflies (Insecta: Symphyta) use their ovipositors to incise plant tissue and deposit eggs, a task that demands precise substrate discrimination while preserving both ovipositor and host. How a passive selective mechanical system such as the ovipositor can discriminate between substrates based on material properties without active sensing or control remained until now an open question. This article presents an analytical model of the selective cutting mechanism inspired by sawfly ovipositors. Substrate cutting is shown to depend on an ultimate stress threshold determined by the interaction between blade geometry and substrate properties. The model captures the transition between cutting and ejection as a competition between a failure and an ejection criterion. Morphological modifiers, including serrulae and banding patterns, are shown to influence the ultimate stress threshold and are strongly dependent on both substrate hardness and elastic modulus. The model explains key previously reported observations obtained from prior experimental work that used scaled up biomimetic blades. These findings suggest novel directions in the design of surgical tools required to cut heterogeneous tissues with minimal extraneous damage.

## 1 Introduction

Tasks requiring interaction with heterogeneous materials, composites or multi-material structures is a challenge across biology, engineering, and medicine. In surgery and soft robotics, for example, tools are needed to operate without damaging critical structures, or to cut, flatten or destroy some whilst preserving others [1–4]. In such cases, precision and control of the intervention based on the structures to be modified or preserved is crucial, especially when the access to sensory input or feedback is limited. In nature, insects have provided several principles which address the need for precision and control in delicate procedures as those used in medical applications. Whether it is mosquito proboscises [5], bee stingers [6,7] or wood-wasp ovipositors [8–10], those principles have proven transferable and useful for multiple applications [11]. This article presents a novel principle based on the ovipositor of the sawfly.

The female sawfly (Insecta: Symphyta) lays her eggs in plants by using her ovipositor to cut a pocket in the plant’s soft tissue [12,13]. The ovipositor comprises two lancets, akin to saw blades, that slide past each other in a reciprocating motion along their longitudinal axis [14]. Each stroke of this motion causes the teeth on opposing blades to trap and mechanically load the substrate from both sides. Depending on the mechanical resistance of the substrate, this interaction results in either substrate fracture or ejection. A selective cutting mechanism (SCM) has been elucidated and modelled in this article. This SCM ensures that the vascular bundles, vital for the survival of the plant are undamaged, protecting thereby the future of the larva. Remarkably, the cutting discrimination is achieved passively as the ovipositor relies on its morphological and mechanical properties as well as those of the substrate that it is meant to interact with. Therefore, the SCM of the sawfly’s ovipositor is an ideal model for the development of bioinspired passive selective cutting tools.

The cutting mechanism was studied experimentally using isometrically scaled up ovipositor blades from two species of sawfly: *Rhogogaster scalaris* (Klug, 1817) and *Hoplocampa brevis* (Klug, 1816) [14]. The study highlighted the existence of an ultimate stress threshold, *σth*, below which substrates are damaged. The research also unveiled a synergistic effect that enables cutting of materials with higher ultimate stress, caused by two morphological traits of the teeth: the serrulae, which are saw-like serrations on the saw teeth, and the anterior tooth, hereafter referred to as bump, which is a protrusion on the basal side of the saw teeth (Figure 1). The research article did not explain however the underlying physical mechanisms, nor did it provide a way to exploit the mechanism. This article addresses this gap, by developing an analytical model of the SCM to explain and predict the behaviour of the cutting mechanism. The two possible outcomes, ejection and cutting, are analytically framed and compared to assess which condition is met first. A force equilibrium is evaluated for the ejection condition and the stress state in combination with material failure criteria for the cutting condition. Two stress formulations are introduced to accommodate different levels of available substrate geometry.

**Figure 1:**
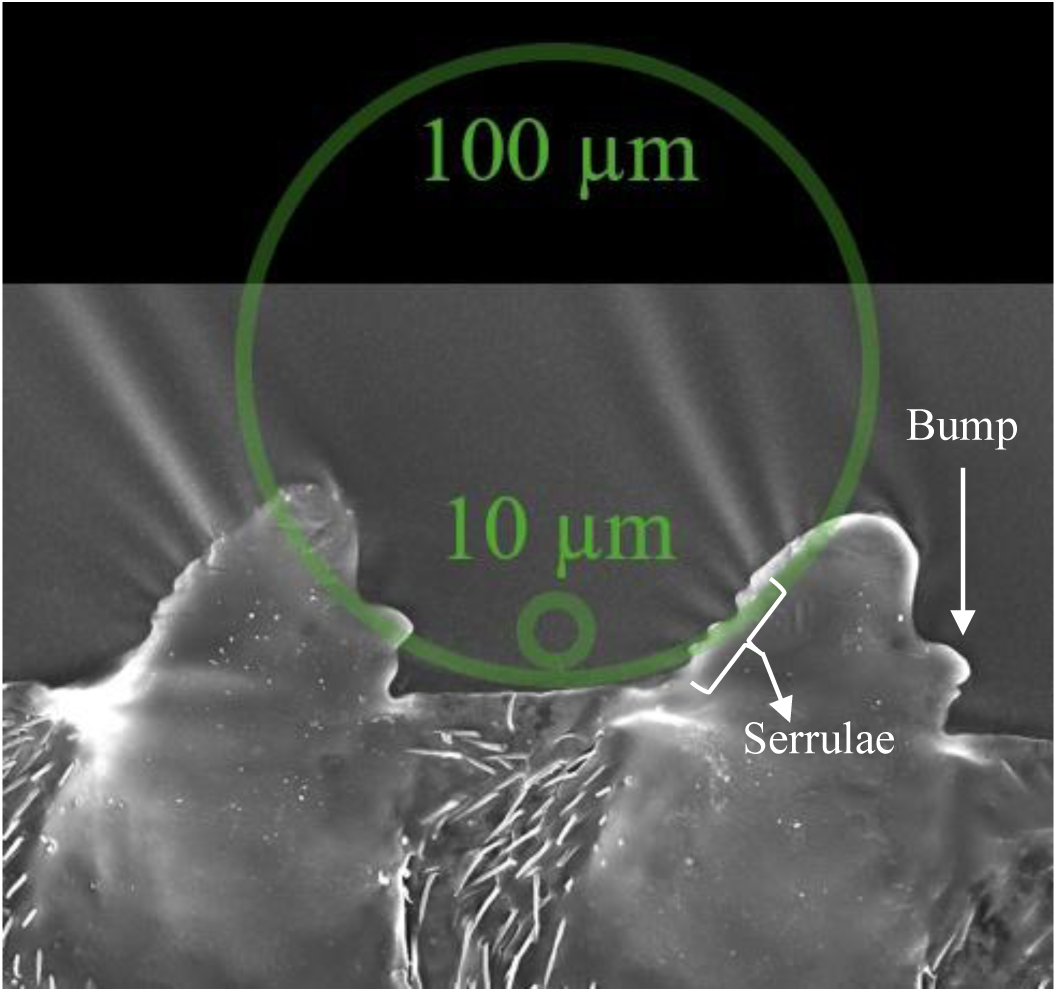
Relative scale of plant cells and R. scalaris ovipositor teeth (8th and 9th). Adapted from [14]. Open access (CC-BY 4.0).

The model, validated here against experimental data from a related study [14], successfully predicts the ejection or damaging outcomes of different blade-substrate combinations from the experimental study. Blade types included serrulae (S), bump (B), and bump–serrulae (BS) designs, tested across agar substrates of 3, 5, and 10 wt%, and 10 wt% ballistic gelatine. The effect of scaling the BS blade is also successfully predicted. In addition, the model can also account for two novel modifiers: the serrulae modifier and the banding pattern modifier which introduce substrate hardness and relative elastic modulus sensitivity, as critical factors to account to during cutting.

The paper is laid out as follows: Section 2 presents the theoretical framework for the selective cutting mechanism, including the approach taken, and the ejection and failure criteria. Section 3 validates the model against experimental data obtained in a previously submitted study. Section 4 explores the effects of some morphological traits of the ovipositor and introduces the effect of the modifiers on the selective cutting mechanism. Section 5 discusses the limitations of the model and possible directions for future work. Section 6 concludes with a summary of the findings along their implications for the design of bioinspired selective cutting tools.

## 2 Theoretical framework

### 2.1 Approach

A tooth in the middle section of an ovipositor of a *R. scalaris* is approximately 34 µm tall. Plant cells are commonly between 10 µm and 100 µm in diameter [15]. The relative sizes of cells and teeth are illustrated in Figure 1 and suggest that the ovipositor is cutting plant tissue at a cellular level. Hence, the analytical model considers a plant cell embedded within the plant tissue as the representative substrate as highlighted in black in Figure 2. Since in the experiments, the system remains in equilibrium under slow loading until either cutting or ejection occurs, a quasistatic modelling approach is appropriate [14].

**Figure 2:**
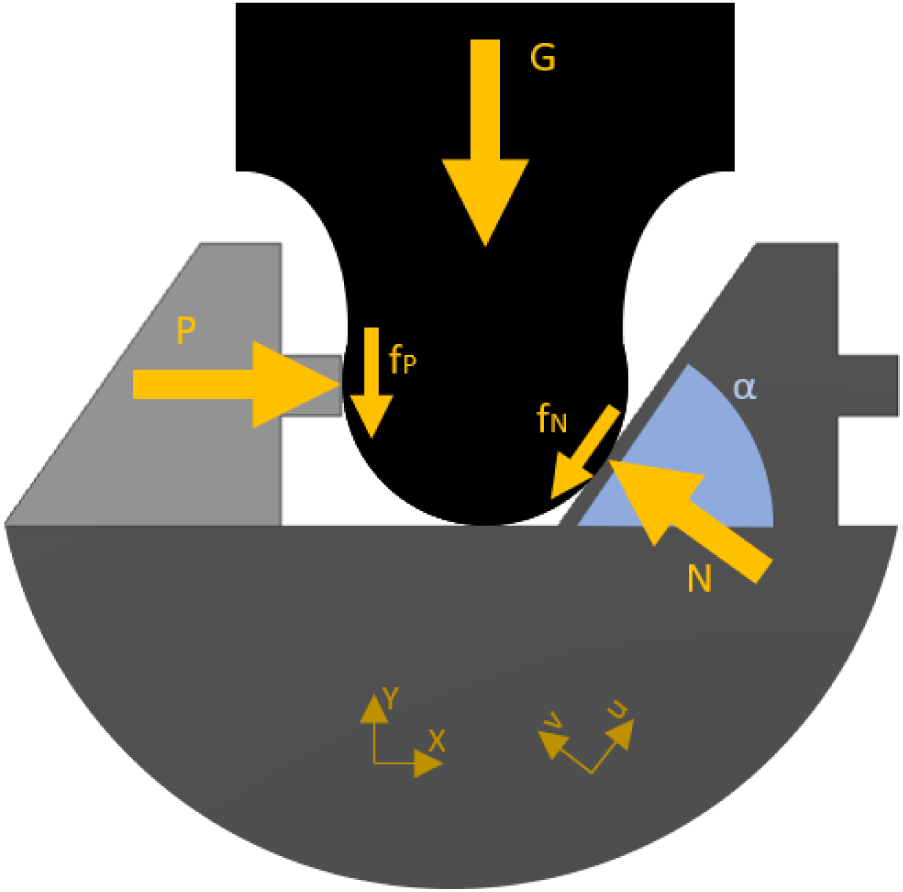
Forces considered in the model. P= cut force, G = force against the substrate, fP = friction force on the bump, fN = friction force on the inclined plane of the tooth, α = tooth angle.

Although the model is adjustable, the variables used in our model are based on data obtained from our experiments [14]. The contact areas and tooth angles have been obtained from the computer-assisted design (CAD) model of the blades modelled using the 9^th^ tooth of the ovipositor of *R. scalaris* (Figure 1). The friction coefficient between the gels and the PMMA blades in the absence of serrulae has been estimated at 0.04 as gels typically exhibit values of friction coefficient ranging between 0.1 and 0.001 [16–18]. When the serrulae are present, they can pierce and embed themselves in the substrate. In such cases, the force opposing the movement, *fN*, as shown in Figure 2, would be more akin to the force necessary to saw the substrate using the serrulae as a saw. Although there are ways to approach the calculation of such forces, including them is not essential to the current level of model complexity [19,20]. Given the purpose of the model in predicting the behaviour of the system, the effect of the serrulae is approximated using a value of the friction coefficient set at 0.35.

### 2.2 Equilibrium of forces

A plant cell can be cut during the reciprocating motion by two teeth, one from each blade. As shown in Figure 2, the two primary forces are the force, *G*, pushing the substrate against the blades and the cut force, *P*, which slides the two blades in opposite directions, trapping the substrate between the two teeth. These forces generate two frictional forces, one, *f_N_*, on the inclined plane of the basal tooth and one, *f*_*P*_, on the bump of the apical tooth. Figure 2 shows also the tooth angle and the two coordinate systems used in the calculations.

During the cutting, a substrate can experience different states, as schematically represented in Figure 3. Initially (label A), the substrate is within the teeth range and is undamaged. As the cutting motion proceeds, two outcomes are possible. Either the substrate is ejected without damage (label B), or it breaks without being ejected (label C). The simultaneous ejection and failure of the substrate (label D) is not physically possible. For the substrate to break, the force required for ejection cannot be delivered and vice versa. The schematics in Figure 3 are used to represent these states throughout the analysis. The model predicts whether the substrate is cut or pushed out of the range of the teeth, based on whether failure conditions or ejection conditions are met.

**Figure 3:**
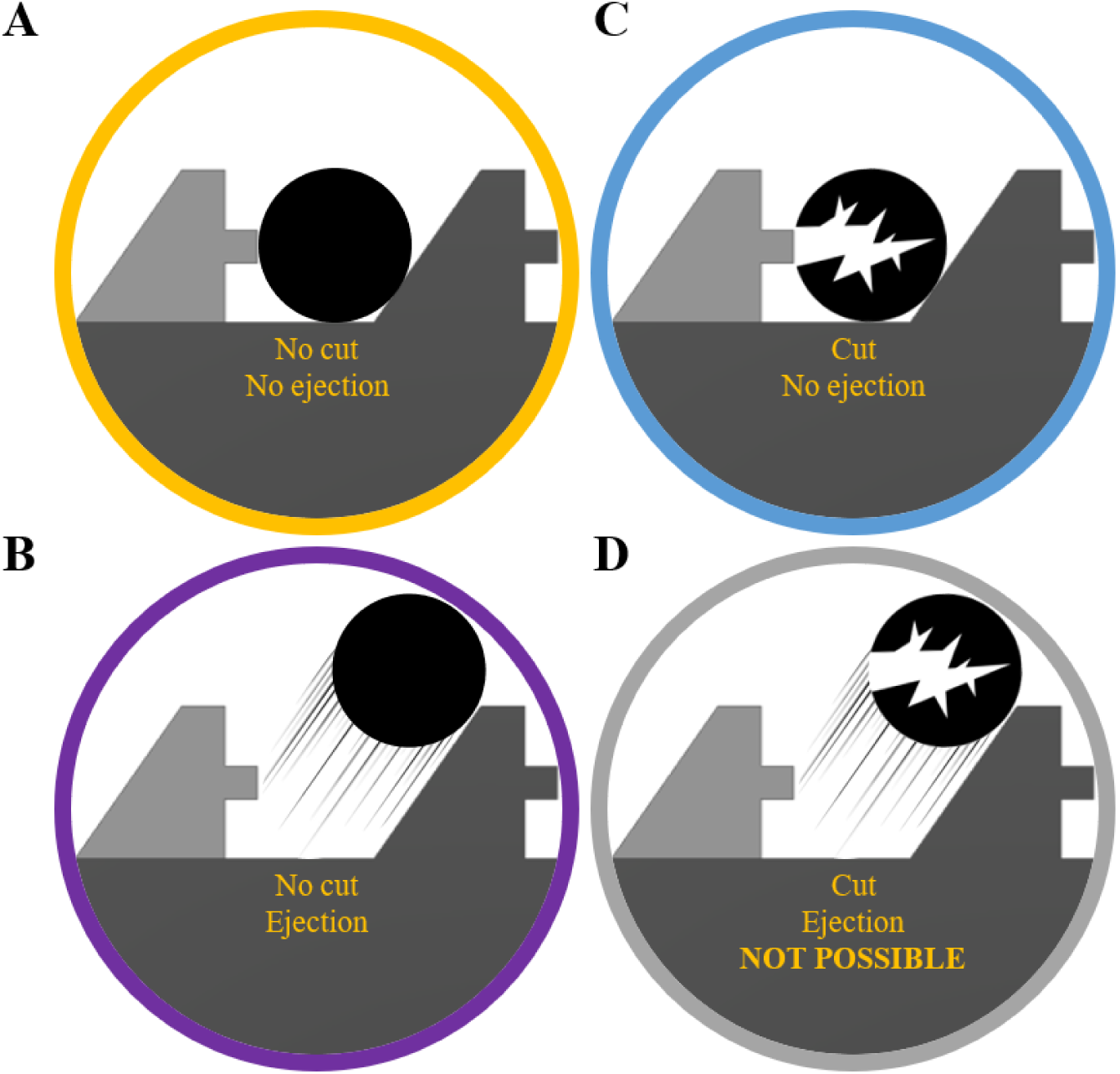
Schematics of model scenarios.

### 2.3 Ejection criterion

For the system to be in equilibrium, the component of the cutting force along the *u-*axis and the *v-*axis, as shown in Figure 2, must equal the sum of opposing forces along the each axis:

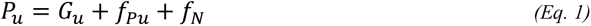

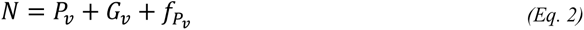

Using trigonometry, and defining two friction coefficients *m* and *n*, such that *f*_*P*_ = *m P*

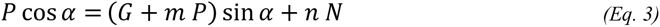

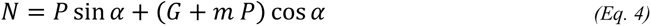

and *f_N_* = *n N*, yields:

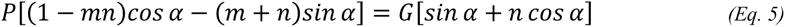

Solving this system to eliminate *N*, the relationship between *P* and *G* is obtained:

Where *m* is the friction coefficient on the bump surface and *n* is the friction coefficient on the tooth inclined plane. This expression defines the relationship between *P* and *G* for specific values of *m*, *n,* and *α*, as represented by the green line in Figure 4. The substrate is ejected without damage when the values of *P* and *G* are below this line, ensuring the integrity of the blade. For the substrate to remain within teeth range and be damaged, the combination of the two forces must be above the line.

**Figure 4:**
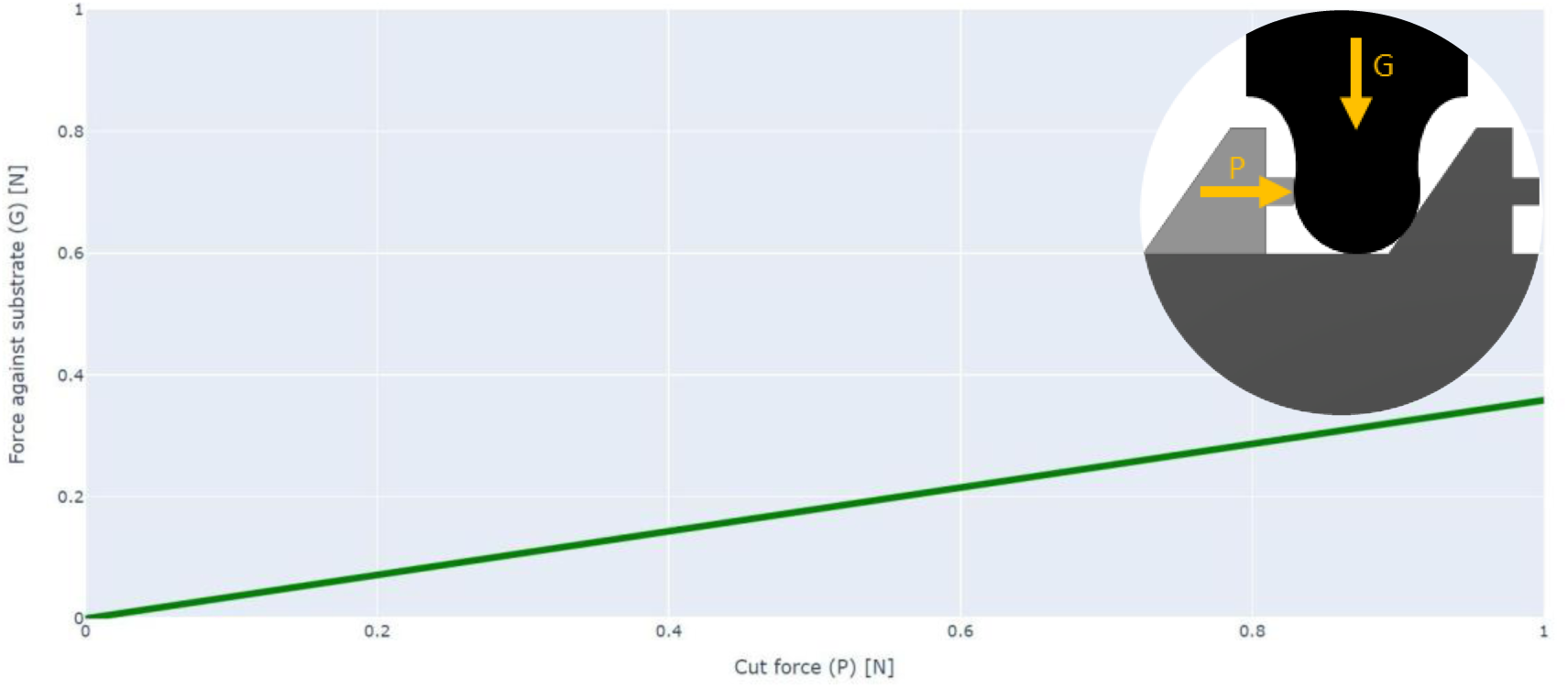
Representation of ejection criteria for m=0.35, n=0.04 and α= 51.57° (0.9 rad). Cut force vs. force against the substrate. Above the line, the substrate remains within teeth range. Underneath the line, the substrate is ejected.

### 2.4 Stresses and failure criteria

The stress state within the substrate is evaluated to establish an appropriate failure criterion. To obtain stress values, the applied forces are first divided by the contact area between the substrate and the teeth. The stresses are distributed along two coordinate systems, the *u-v-w* and the *x-y-z*, as shown in Figure 5. The friction force on the tooth inclined plane, *f_N_*, generates a shear stress acting on the plane normal to axis *v* in the direction of axis u, τ_*vu*_, as shown in Figure 5. Since the substrate is a cell embedded in plant tissue and prevented from rotating, an opposing stress of equal magnitude, τ_*vu*_, appears.

**Figure 5:**
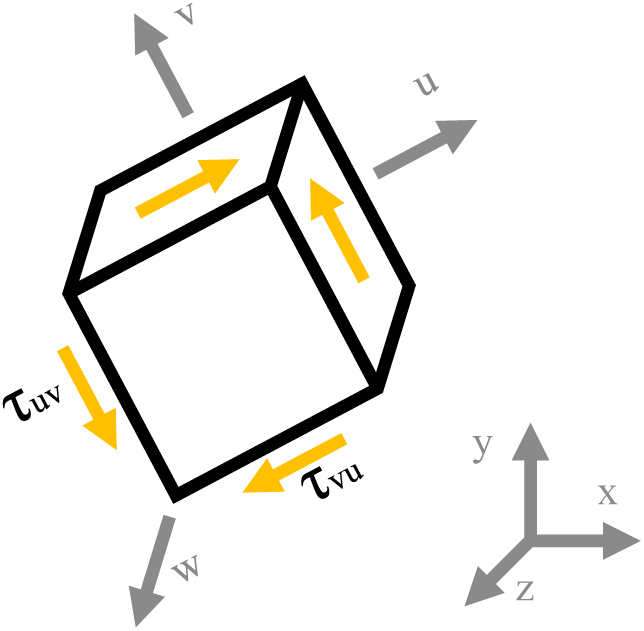
Schematics of stresses aligned with axis u-v-w.

The stress tensor *U*_*vuw*_ along the *u-v-w* coordinate system is given by (Eq. 6):

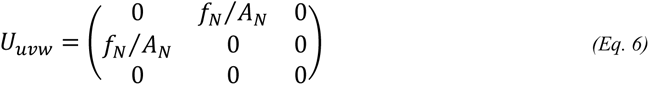

Where *A*_*N*_ is the contact area between the teeth inclined plane and the substrate. By using a rotation matrix around the out-of-plane axis *w*, the stress tensor *U*_*vuw*_can be aligned with the coordinate system *x-y-z* to obtain the *U*′_*xyz*_ stress tensor.

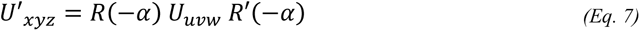

Where *R* is the counterclockwise rotation matrix, *R*′ is the transposed rotation matrix. Two modelling approaches are considered to evaluate the stresses aligned with the axis *x-y-z*. In the first one, the stress generated by the cut force is considered as a compressive stress. This reduces the number of variables alongside the required information about the substrate, which simplifies the model at a cost of introducing some inaccuracy. In the second approach, the stress generated by the cut force is considered as a shear stress. While this reflects more accurately the nature of the stress state, it is also less practical because the shear area *A*_*PS*_ is related to the thickness of the substrate which can be variable. Both models have been built and are described here.

#### 2.4.1 Model using compressive stress

When the cut force *P* in the *x-y-z* coordinate system is considered as a compressive load, the corresponding stress is obtained by dividing *P* by the contact area between the bump and the substrate. This stress acts on the plane normal to the *x*-axis in the direction of the same axis σ_*xx*_. On the *y*-axis, the force against the substrate *G* produces a compressive stress σ_*yy*_, as shown in Figure 6, calculated by dividing *G* by the projected area of the inclined plane of the tooth which is normal to the applied force. The friction force on the bump *f*_*P*_generates a shear stress τ_*xy*_ acting on the plane perpendicular to the *x*-axis and directed along the *y*-axis as shown in Figure 6. Since the substrate is not rotating, this stress is accompanied by an equal one in the opposite direction τ_*yx*_.

**Figure 6:**
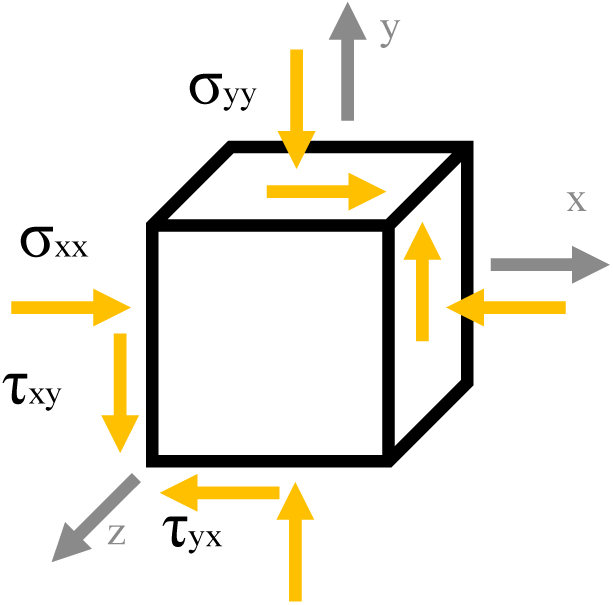
Schematics of stresses aligned with axis x-y-z assuming compressive load.

The resulting stress tensor in the *x-y-z* coordinate system is:

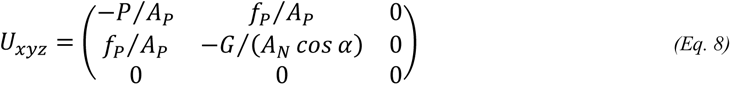

Where *A*_*P*_ is the contact area between the substrate and the bump. The stress tensors *U*′_*xyz*_ and *U*_*xyz*_ are summed to obtain the total stress tensor σ_*xyz*_.

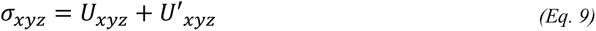

#### 2.4.2 Model using the shear stress

If the cut force *P* in the *x-y-z* coordinate system is considered to generate a shear stress, more geometric information about the substrate is needed. In this case, the area used to compute the stress is defined by the thickness of the substrate in the *x*-direction and the height of the bump contacting the substrate in the *y*-direction. This results in a shear stress τ_*zx*_ and its counterpart τ_*xz*_ since a cell does not rotate within the plant tissue. The force against the substrate *G* continues to generate a compressive stress σ_*yy*_, and the friction force on the bump *f*_*P*_ still produces a shear stress τ_*xy*_, as shown in Figure 7.

**Figure 7:**
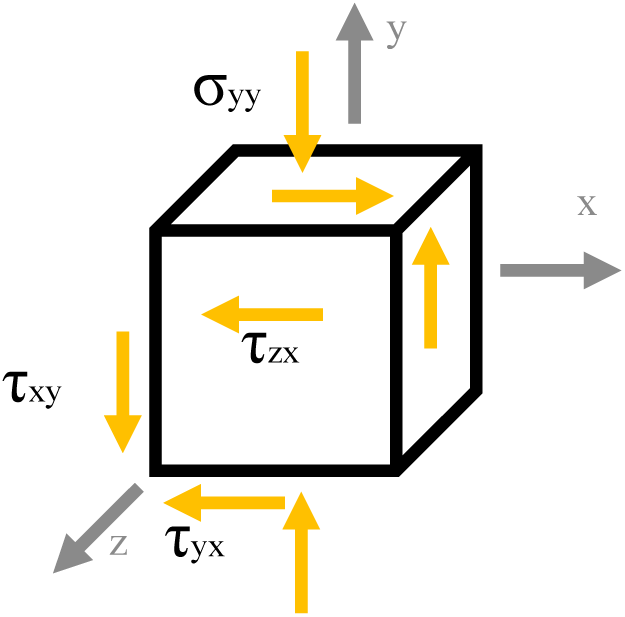
Schematics of stresses aligned with axis x-y-z in the shear case.

The resulting stress tensor in this case is given by:

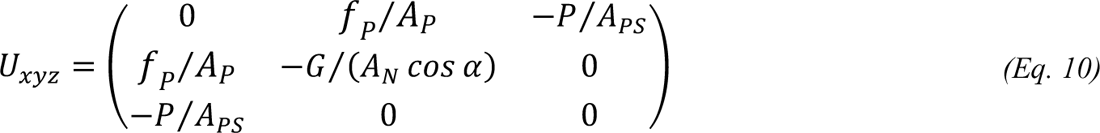

Where *A*_*PS*_is the shear area between the bump and the inclined plane of the tooth. The stress tensors *U*′_*xyz*_ and *U*_*xyz*_ are summed to obtain the total stress tensor σ_*xyz*_.

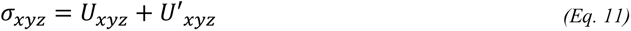

#### 2.4.3 Failure criteria

A computational model was developed to obtain the total stress tensors of the system. It also calculates the principal stresses, the hydrostatic stress tensor, the deviatoric stress tensor, and the failure criteria. Several failure criteria have been implemented. These include von Mises, based on distortion energy [21,22]; Tresca, based on maximum shear stress [22,23]; the maximum normal stress criterion [22,24]; and the modified Mohr criterion [22,25]. The first two are generally applied to ductile materials, while the latter two are typically preferred for brittle materials. Most of these criteria require knowing only the ultimate compressive strength of the substrate. However, the modified Mohr criterion additionally requires the ultimate tensile stress.

Considering plant cells as a ductile or brittle material would be inaccurate due to their highly complex structure and wide variability of their mechanical properties [26]. Nonetheless, the mechanical behaviour of some plant tissues has been simulated with visco-elastoplastic behaviour, with failure prediction using the von Mises effective stress [27]. Although gels are often considered generally brittle, agar is amongst the less brittle gels [28]. The compressive stress tests performed on the agar substrates of our previous experimental work did not show a pronounced brittle behaviour [14]. Von Mises stresses and strains have been successfully used to predict the behaviour of agar in simulations [29–31]. Taking all this into account, the von Mises failure criterion has been selected for this analysis. This criterion evaluates the effect of a complex stress state of a material by calculating an equivalent stress based on the distortion energy. Yielding is predicted when this equivalent stress surpasses yield strength of the material. All the named failure criteria are implemented in the model and can be used to tailor the model to the behaviour of other substrates. In the following analysis, the cutting force is treated as a compressive load.

A three-dimensional plot would illustrate the behaviour of the substrate-ovipositor system. To improve readability, Figure 8 shows a two-dimensional projection with isolines representing combinations of forces which yield the same von Mises effective stress. The substrate will be damaged if the combination of force against the substrate (*G*) and cut force *P* lies beyond the ultimate strength isoline of the substrate.

**Figure 8:**
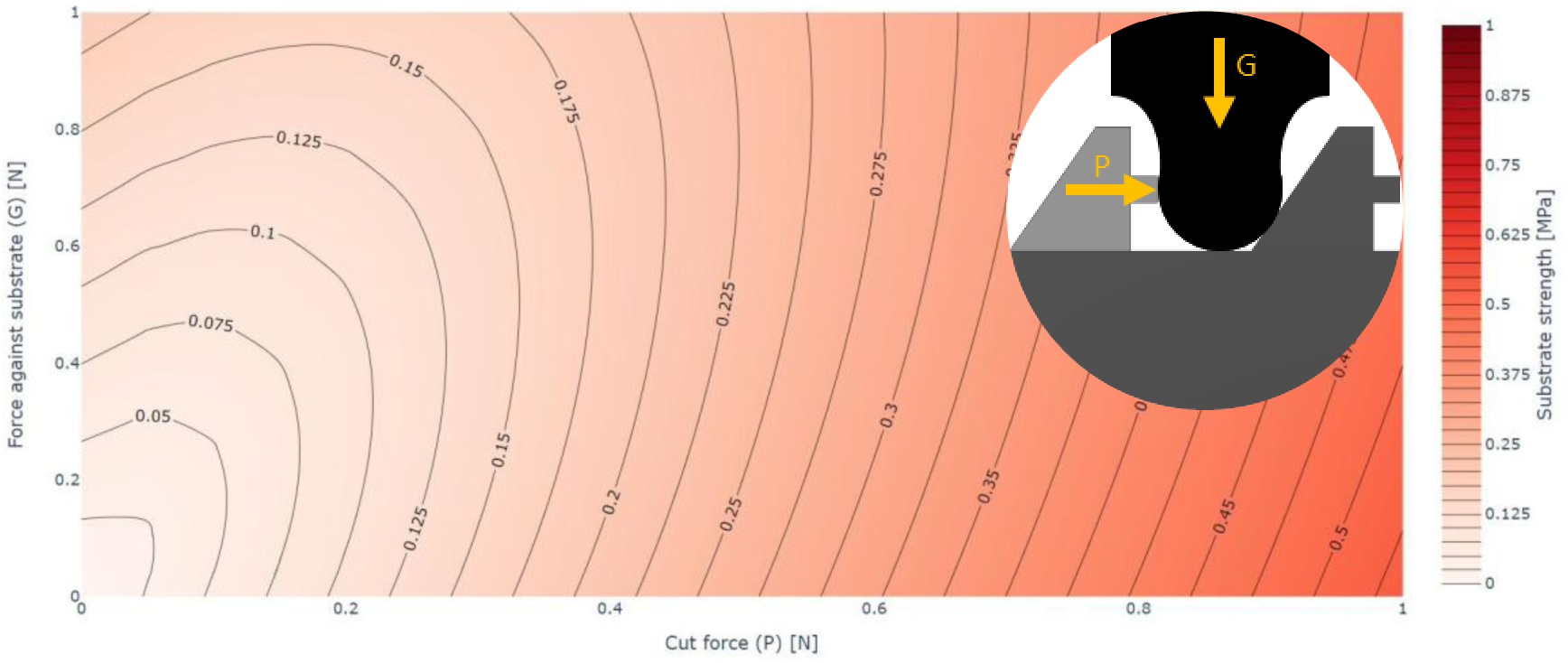
Graph of the relationship between cut force P, force against the substrate G, and the Von Mises stress. Tooth angle α =51.57° (0.9 rad). Bump friction coefficient, m = 0.04. Tooth inclined plane friction coefficient, n = 0.35. Scale = 400. Bump contact area Ap = 1.751 mm^2^. Tooth inclined plane contact area An = 10.030 mm^2^.

### 2.5 Selective cutting mechanism model

The SCM model is obtained by combining the substrate ejection and material failure criteria, as shown in Figure 9.

**Figure 9:**
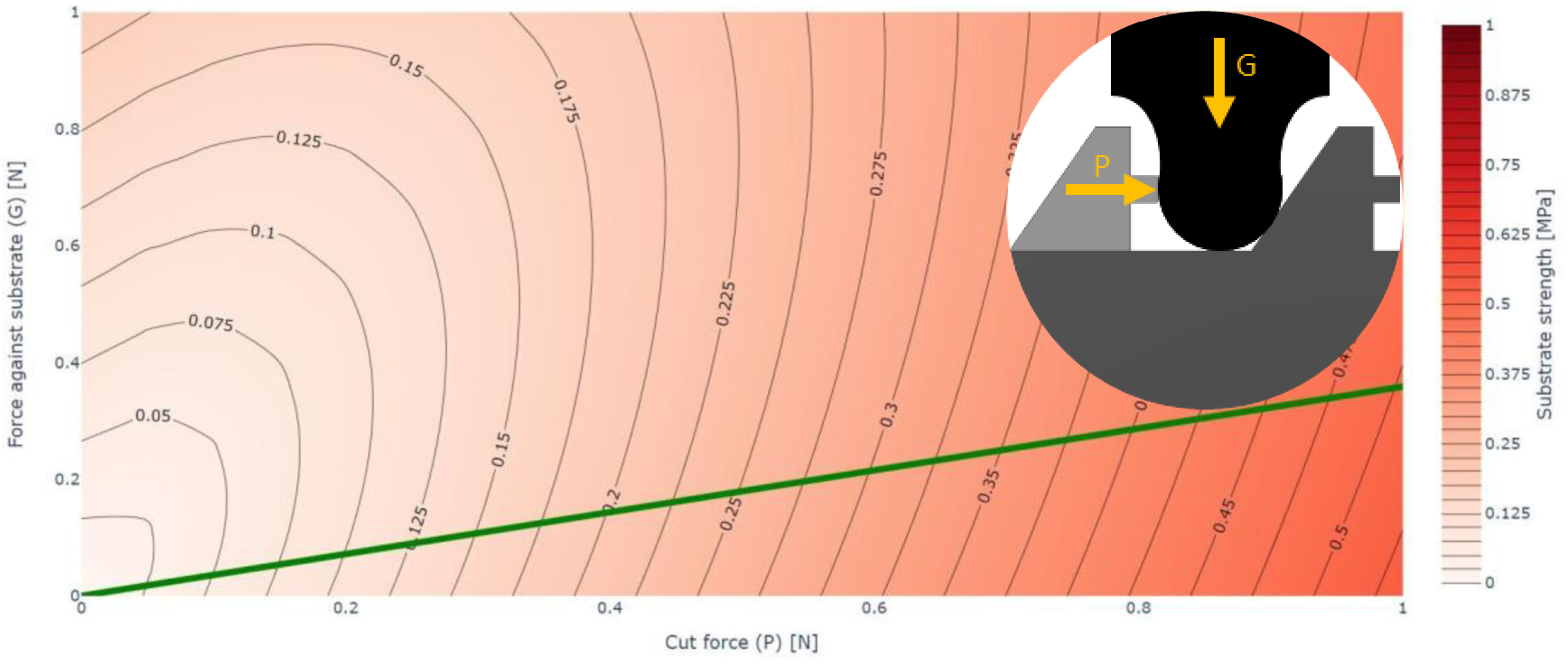
Graph of the SCM showing the relationship between the cut force P, force against the substrate G, and the Von Mises stress. Variables include tooth angle, α = 51.57° (0.9 rad), friction coefficient of the bump, m = 0.04, friction coefficient of tooth inclined plane, n = 0.35. Scale = 400, contact area of the bump, Ap = 1.751 mm^2^, and contact area of the tooth inclined plane, An = 10.03 mm^2^.

Figure 10 predicts the behaviour of a system whose substrate has an ultimate compressive strength of 0.3 MPa, represented by the corresponding isoline. Beyond this isoline, the substrate fails. Below the ejection line, the substrate will be ejected from the teeth range. The failure isoline and ejection line delimit therefore four quadrants whereby the combination of cut force and force against the substrate causes different behaviours of the system as shown in Figure 10. These regions explain the SCM previously observed in our experimental study [14].

**Figure 10:**
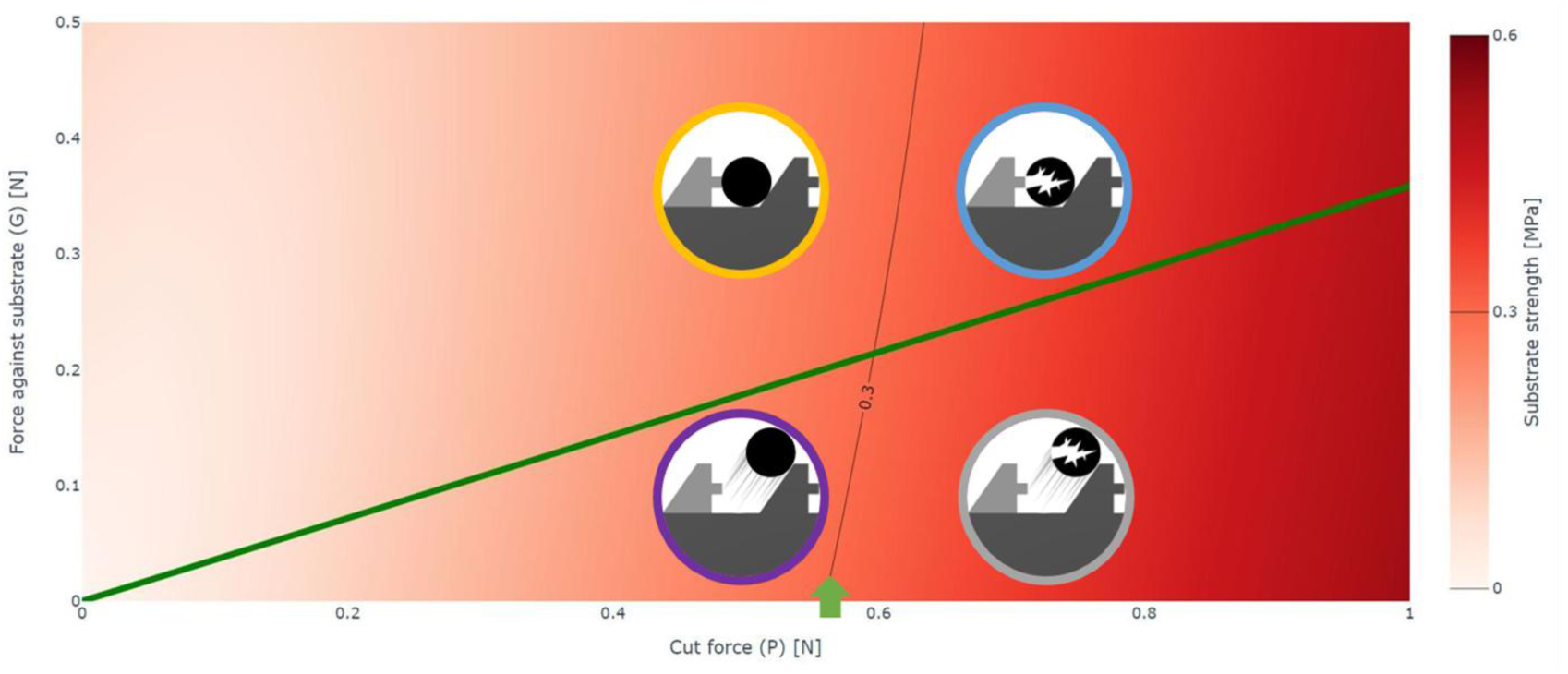
Four SCM model regions based on combination of forces: top left is stable with no damage or ejection, top right involves damage without ejection, bottom left leads to ejection without damage, and bottom right represents a physically impossible case of simultaneous damage and ejection.

When the substrate cannot be cut, it is ejected, preserving the mechanical integrity of the teeth. Given a large enough cut force *P*, a substrate with an ultimate strength of 0.3 MPa will be ejected without damage as long as the force against the substrate remains below 0.2 N. With a force against the substrate higher than 0.2 N the substrate is not ejected and fails. Similarly, if the force against the substrate remains at 0.2 N but the ultimate strength of the substrate decreases to 0.25 MPa, the substrate is damaged.

## 3 Experimental validation of the model

The model was validated against the experimental data obtained using isometrically scaled blades based on the ovipositor of *R. scalaris*, as described in our experimental study [14]. Although the cutting experiments did not constrain substrate rotation, unlike the model, the comparison remains valid since none was observed during testing. This discrepancy might have introduced minor inaccuracies, but these are considered negligible for the current analysis. The contact areas and the force against the substrate, *G*, were taken from the experimental study. The model parameters *α*, *n* and *m* were set as explained in Section 2.1, and the cut force *P* was calculated accordingly. Since the experimental rig provided effectively unlimited cut force, any substrate should be damaged if the force against the substrate is high enough [14]. A summary of all experimental test points, model variables, calculated cut force *P*, and cutting outcomes is presented in Table 1.

**Table 1:** Experimental outcomes across blade types, scales, and substrate conditions. Each row corresponds to a single test point, showing the blade configuration, material properties, and whether substrate damage occurred (“Exp. damage”). Shaded cells indicate cases where no cutting was observed. Blade abbreviations: BS = blade with bump and serrulae; B = blade with only bump; S = blade with only serrulae. Variables: AP = bump contact area, AN = inclined plane contact area, α = tooth angle, n = inclined plane friction coefficient, m = bump friction coefficient, σcu = substrate’s compressive failure stress, G = applied force against the substrate, P = predicted cutting force.

| Blade | Scale | $A_P$ [mm<sup>2</sup>] | $A_N$ [mm<sup>2</sup>] | $\alpha$ [rad] | $n$ | $m$ | Point | Substrate | $\sigma_{cu}$ [MPa] | $G$ [N] | $P$ [N] | Exp. damage |
| --- | --- | --- | --- | --- | --- | --- | --- | --- | --- | --- | --- | --- |
| BS | 400 | 1.751 | 10.03 | 0.9 | 0.35 | 0.04 | A | 3 wt% agar | 0.082 | 0.148 | 0.177 | Yes |
|  |  |  |  |  |  |  | B | 5 wt% agar | 0.115 | 0.148 | 0.242 | Yes |
|  |  |  |  |  |  |  | C | 10 wt% ballistic gelatine | 0.18 | 0.148 | 0.366 | Yes |
|  |  |  |  |  |  |  | D | 10 wt% agar | 0.3 | 0.148 | 0.593 | No |
|  |  |  |  |  |  |  | E | 10 wt% agar | 0.3 | 0.216* | 0.605 | Yes |
| BS | 200 | 0.438 | 2.905 | 0.9 | 0.35 | 0.04 | F | 3 wt% agar | 0.082 | 0.148 | 0 | Yes |
|  |  |  |  |  |  |  | G | 5 wt% agar | 0.115 | 0.148 | 0.059 | Yes |
|  |  |  |  |  |  |  | H | 10 wt% ballistic gelatine | 0.18 | 0.148 | 0.101 | Yes |
|  |  |  |  |  |  |  | I | 10 wt% agar | 0.3 | 0.148 | 0.164 | Yes |
| B | 400 | 1.751 | 10.03 | 0.9 | 0.04 | 0.04 | J | 3 wt% agar | 0.082 | 0.148 | 0.162 | Yes |
|  |  |  |  |  |  |  | K | 5 wt% agar | 0.115 | 0.148 | 0.221 | No |
|  |  |  |  |  |  |  | L | 10 wt% ballistic gelatine | 0.18 | 0.148 | 0.337 | No |
|  |  |  |  |  |  |  | M | 10 wt% agar | 0.3 | 0.148 | 0.55 | No |
| S | 400 | 3.36 | 10.03 | 0.9 | 0.35 | 0.04 | N | 3 wt% Agar | 0.082 | 0.148 | 0.359 | Yes |
|  |  |  |  |  |  |  | O | 5 wt% Agar | 0.115 | 0.148 | 0.492 | No |
|  |  |  |  |  |  |  | P | 10 wt% ballistic gelatine | 0.18 | 0.148 | 0.749 | No |
|  |  |  |  |  |  |  | Q | 10 wt% Agar | 0.3 | 0.148 | 1.216 | No |

\* The increased force $G$ is due to an added weight of 7g to the substrate.

As shown in Figure 11 through Figure 14, in all tested cases the model successfully predicts whether a substrate is damaged or ejected across all blade types and substrate conditions. Each point is plotted relative to the ejection line and failure criterion. As explained in Section 2.5, cutting occurs only when the combination of applied forces lies above the ejection line and beyond the failure isoline. In Figure 11, points A to D share an *σth* of approximately 0.25 MPa. Points A to C are located above the ejection line and beyond their respective failure isolines, consistent with observed substrate damage. Conversely, point D lies below the ejection line, meaning that the substrate is ejected. Point E demonstrates how increasing the force against the substrate *G* increases the *σth* to around 0.35 MPa enabling failure of the same substrate which was ejected at point D. The model also predicts that a decrease in scale increases the *σth* for a given *G*, allowing for all the substrates to be cut. This is shown in Figure 12, where all the points lie well above the ejection line.

**Figure 11:**
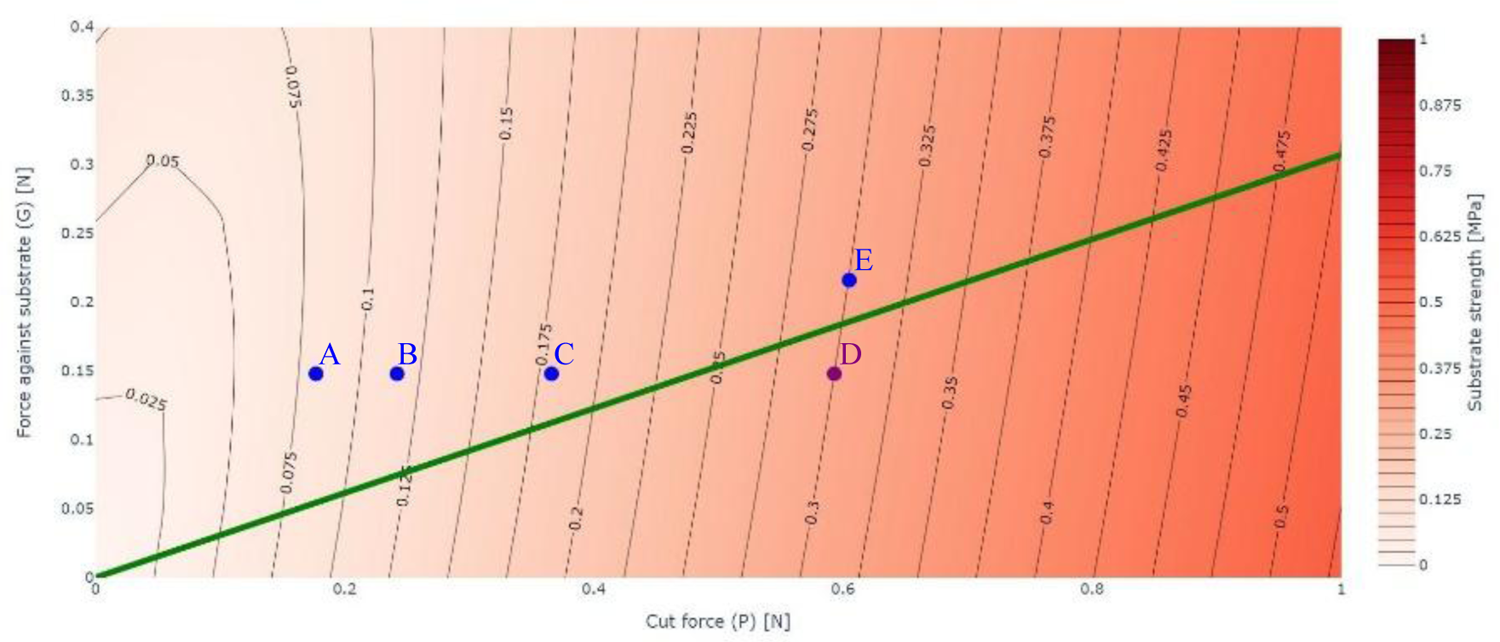
Model predicting ejection-failure outcome of experimental points.The threshold and the effect of adding weight are shown. The information for the points is listed in Table 1.

**Figure 12:**
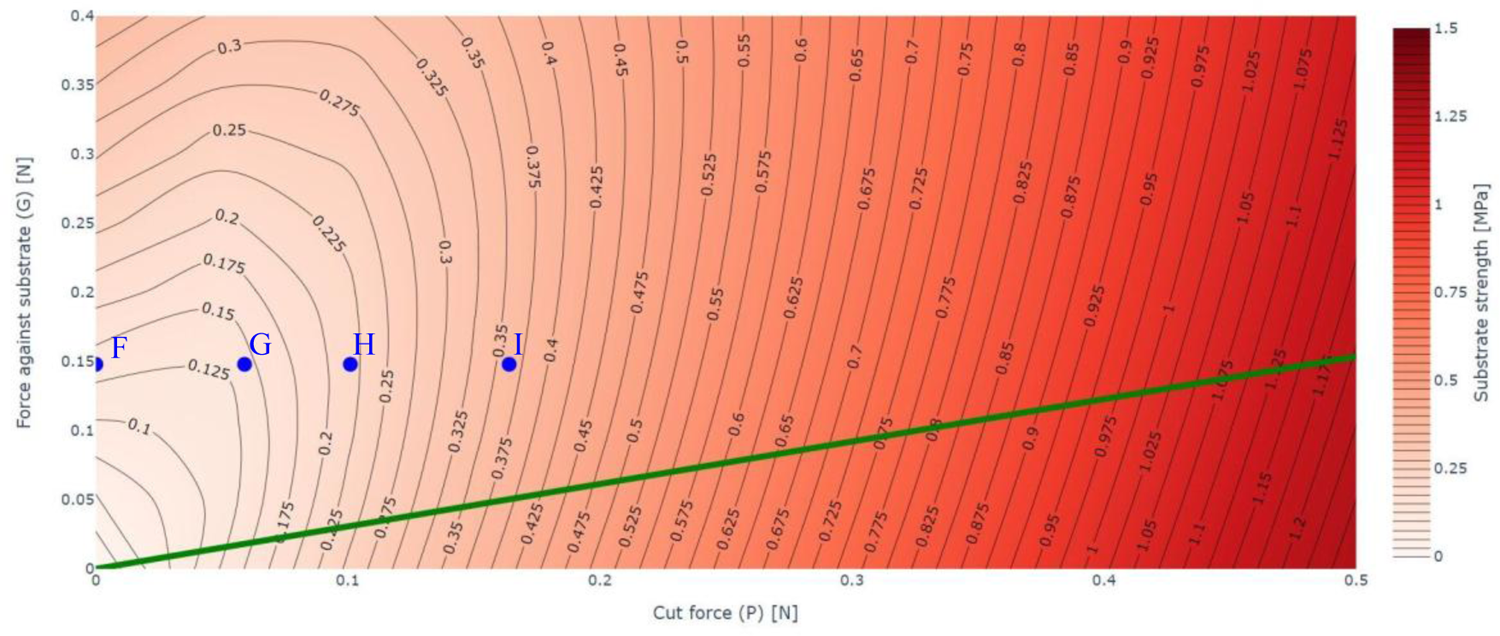
Model predicting ejection-failure outcome of experimental points. The effect of reducing the scale is shown. The information for the points is listed in Table 1. The cut force P for point F is 0 since pushing such a substrate against a blade with a force G of 0.148 N is enough to cut the substrate at that scale.

Figure 13 and Figure 14 demonstrate how removing key blade morphological traits reduces cutting performance. Removing the serrulae or the bump from the blades reduces the threshold, causing the ejection of all the substrates except for the 3 wt% agar. The lack of serrulae is represented by a lower friction coefficient *n* in the tooth inclined plane (Figure 13). An increase to the bump contact area *A*_*P*_represents how, when a bump is lacking, its task is performed by the main body of the tooth (Figure 14).

**Figure 13:**
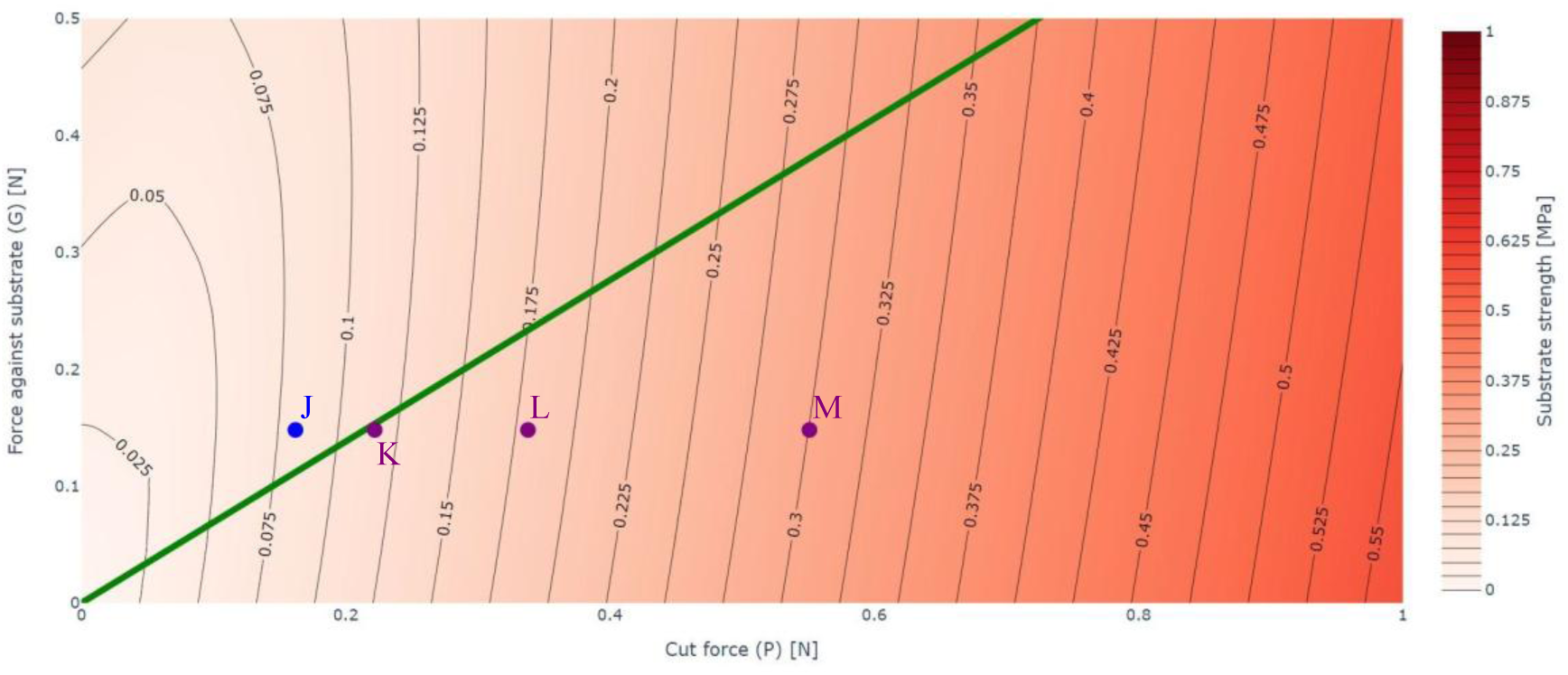
Model predicting ejection-failure outcome of experimental points. The effect of removing the serrulae is shown. Point data listed in Table 1.

**Figure 14:**
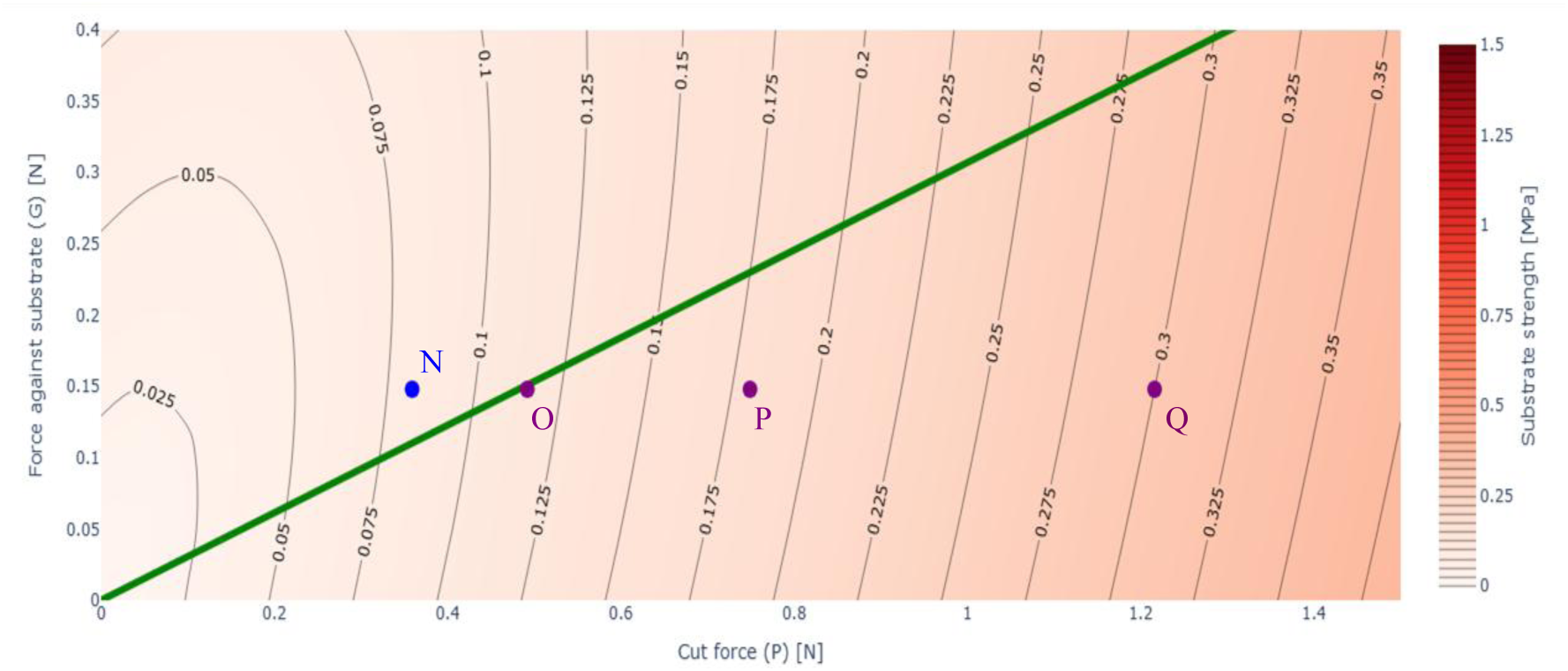
Model predicting ejection-failure outcome of experimental points. The effect of removing the bump is shown. Point data listed in Table 1.

When the cut force *P* is modelled as generating a shear stress, the calculated values necessary to cut each substrate are higher than when the force is assumed to generate a compressive stress. This difference is due to the thickness of the substrate leading to a larger shear area, which requires a greater force to achieve the same stress. On the other hand, the von Mises failure criterion predicts increasingly larger equivalent stresses for shear stresses than for compressive stresses of the same magnitude. This tends to compensate for the difference caused by the area, causing the two models to converge as the cut force increases. The trend continues until they predict the same cut force. Beyond this point, the shear-stress model predicts lower cut forces than the compressive-stress model. This effect is intrinsically related to the von Mises failure criterion; other failure criteria may produce different trends. The maximum principal stress criterion, for example, predicts the same failure stress for the same values of compressive stress and shear stress. In that case, the difference in the predicted cut force is exclusively related to the difference in stress area.

## 4 Selective cutting mechanism modifiers

The model can be regarded as a first-order approximation that accounts for the primary mechanisms involved in the cutting of the plant by the ovipositor. This section presents additional traits that modify the predicted outcomes of the model. The model can be accommodated to account for two morphological modifiers that influence cutting performance: the presence of serrulae and the angle of the teeth.

The serrulae piercing the substrate increases the value of the friction coefficient. The tooth angle modifies the range of substrates that can be cut. The presence of a banding pattern, shown in Figure 15, in some ovipositors suggests that the tooth angle could be modified dynamically during cutting due to the thinner profile and lower elastic modulus of the pattern.

**Figure 15:**
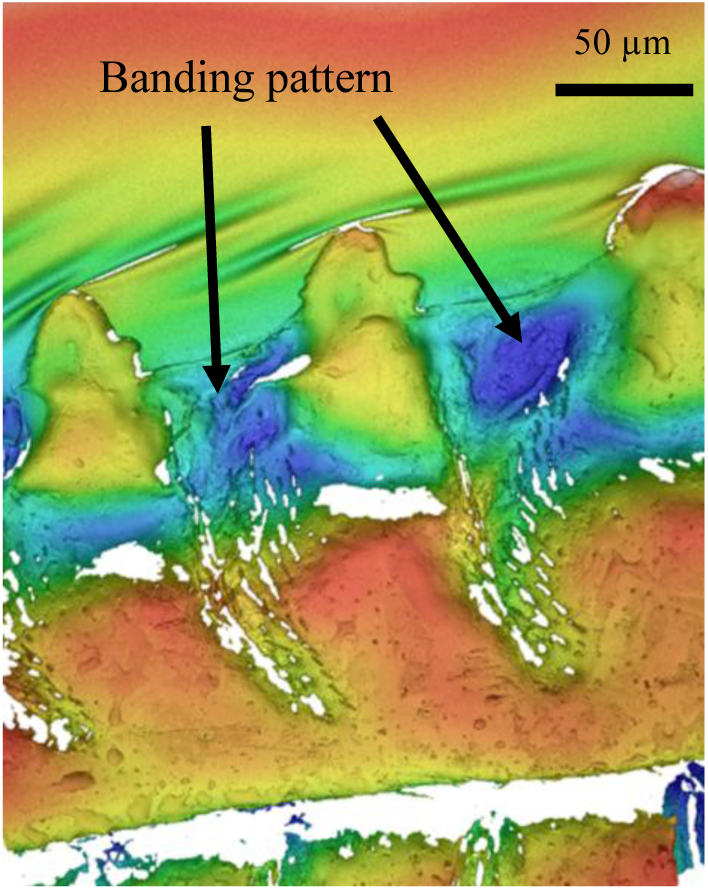
Profilometry of R. scalaris ovipositor showing banding pattern. Adapted from [14]. Open access (CC-BY 4.0).

The banding pattern, further described in Section 4.2 consists of bands between teeth that extend from the edge of the blade to the ramus, a reinforced longitudinal region along the opposite side of the blade. A schematic representation of these two modifiers is provided in Figure 16.

**Figure 16:**
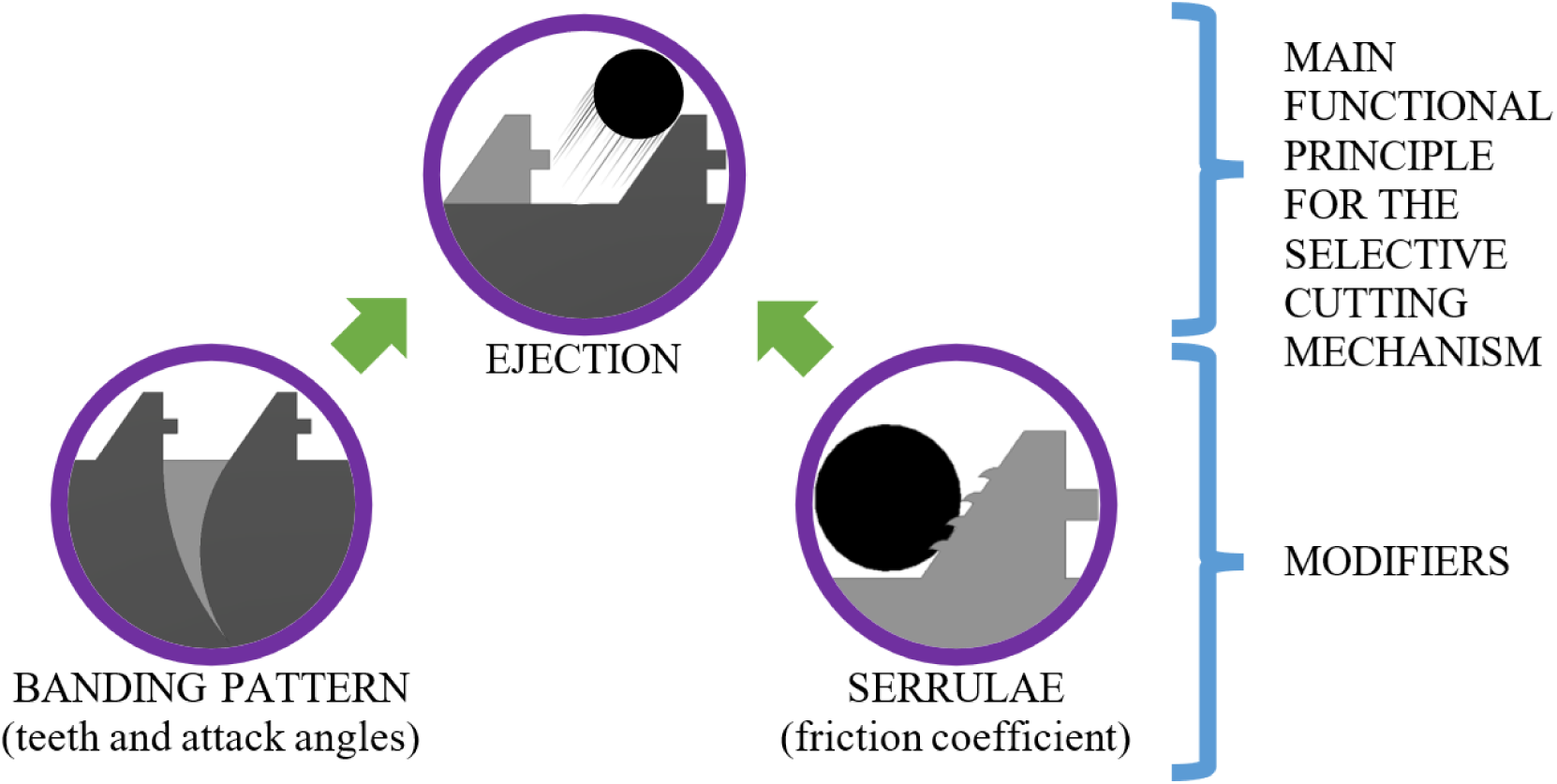
Functional schematics of the modifiers regarding the SCM.

The morphological data used to characterize these features were obtained from a combination of previously published and newly collected imaging. SEM and profilometry images were sourced from a related experimental study [14], with tooth angles measured using ImageJ. The banding patterns were visualized using the profilometry and new confocal laser scanning microscopy (CLSM). The imaging was performed on a Leica SP8 system. A HC PL APO 20x/0.75 CS2 air objective (Leica 506517) was used. The excitation was provided by a Supercontinuum White Light Laser except for when the excitation was at 405 nm which was provided by a Leica SP8 405 nm laser. All detections were made using a Leica HyD hybrid detector. The system was operated by the commercial software Leica Application Suite X (LAS X). The carrier slides used were 25 mm diameter No. 1.5H high precision glass coverslips (Marienfeld Superior 0117650) and in an Attofluor Cell Chamber (Thermo Fisher A7816). The settings used on the CLSM can be found in Table 2. Image stacks were processed into maximum intensity Z-projections using ImageJ.

**Table 2:** CLSM settings.

|  | *R. scalaris* |
| --- | --- |
| **Channel 1 Excitation** | 405 nm |
| **Channel 1 Detection** | 415-475 nm |
| **Channel 2 Excitation** | 488 nm |
| **Channel 2 Detection** | 500-560 nm |
| **Channel 3 Excitation** | 600 nm |
| **Channel 3 Detection** | 650-785 nm |
| **Pixel size** | 94 nm |
| **Z-step** | 562 nm |
| **Pinhole** | 0.75AU (42.5 $\mu\text{m}$) |
| **Magnification** | 20 |

### 4.1 Serrulae as a modifier

Experimental results reported in our companion study [14], together with model predictions, show that reducing the friction coefficient or removing the serrulae significantly lowers the *σth*. Figure 17 plots the friction coefficient of the inclined plane in relation to the force against the substrate and the failure criterion. The cut force *P* was fixed at 0.232 N to plot points B and K from Table 1, it shows that a significant increase in the friction coefficient can disable the ejection mechanism. Values of friction coefficient larger than 0.72 suppress the ejection mechanism entirely.

**Figure 17:**
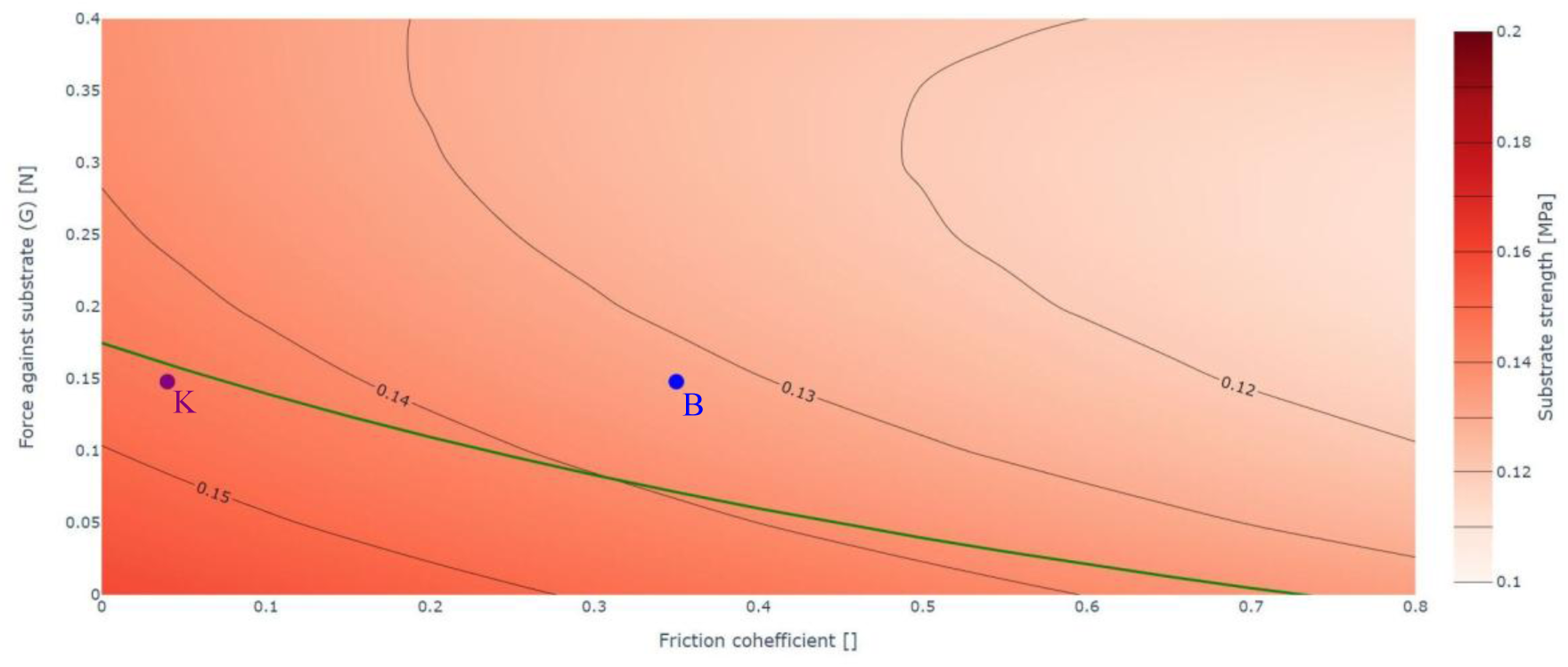
Effect of friction coefficient on ejection threshold. Point data listed in Table 1.

A high value of the friction coefficient may result from the serrulae piercing and hooking into the substrate, as shown in Figure 18, suggesting that the serrulae effect functions as a modifier of the SCM.

**Figure 18:**
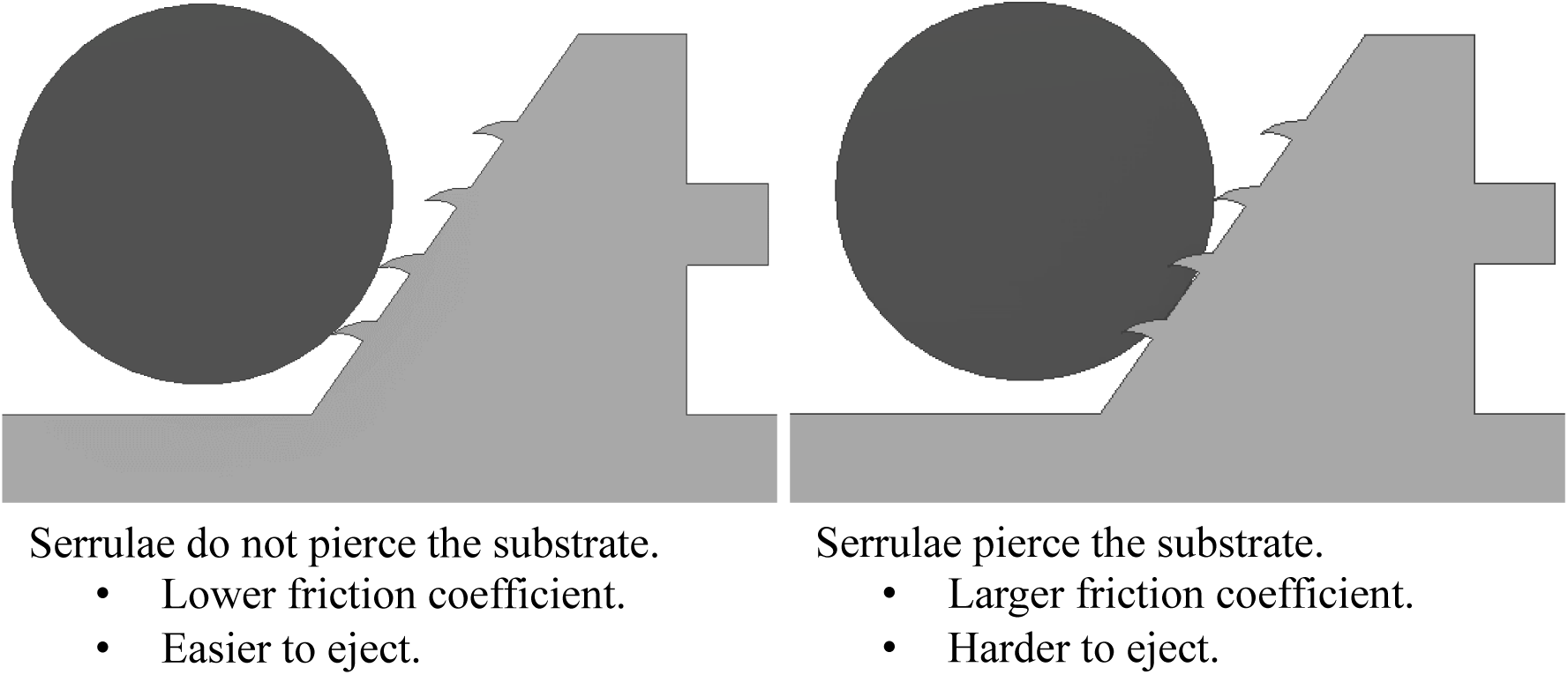
Schematic of serrulae-substrate interaction. Left: low-friction, non-piercing contact. Right: high-friction due to serrulae penetration.

### 4.2 Teeth angle and banding pattern modifier

The model also indicates that the angle of the teeth significantly influences the SCM. This is illustrated in Figure 19 where the tooth angle is plotted in relation to the force against the substrate and the failure criterion for a constant *P* of 0.232 N. Point B lies above the ejection line, indicating substrate failure, in agreement with experiments. For a given ultimate stress, adjusting the tooth angle can either deactivate the ejection mechanism or allow the ejection and therefore preserve the integrity of the blade. As the tooth angle approaches 90° (1.57 rad), the system can cut substrates with higher ultimate strength.

**Figure 19:**
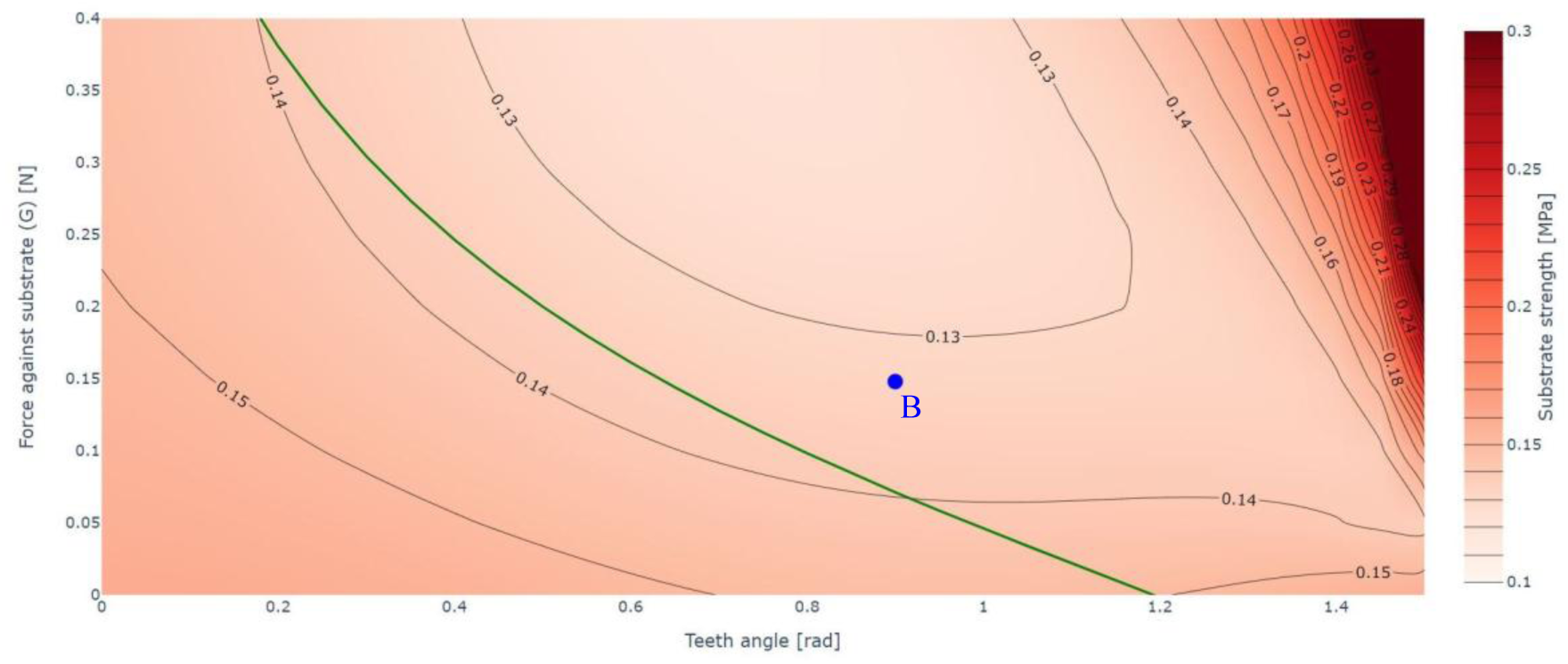
Effect of tooth angle on ejection threshold. Point data listed in Table 1.

The angle of each tooth along the ovipositor has been measured relative to the base of the tooth. These angles are shown in Figure 20. Three distinct regions are colour-coded in Figure 20 and Figure 21 corresponding to different functional roles of the teeth.

**Figure 20:**
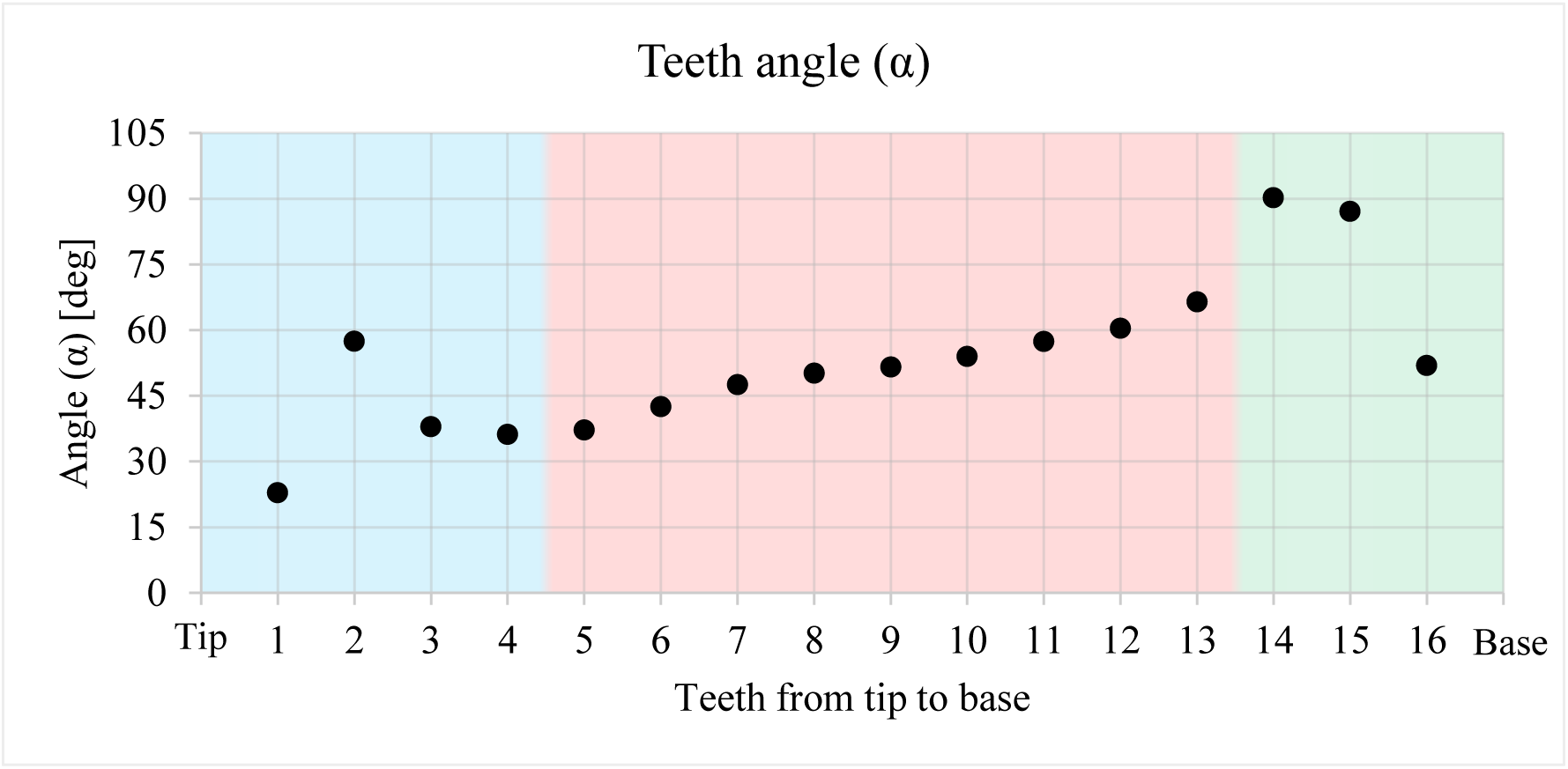
Tooth angle along ovipositor length. The coloured areas are related to different tasks of the teeth. Blue: teeth related to insertion. Red: teeth related to cutting through cells using the SCM to differentiate tissues. Green: tooth related to cutting through the outer stronger layers of cells.

**Figure 21:**
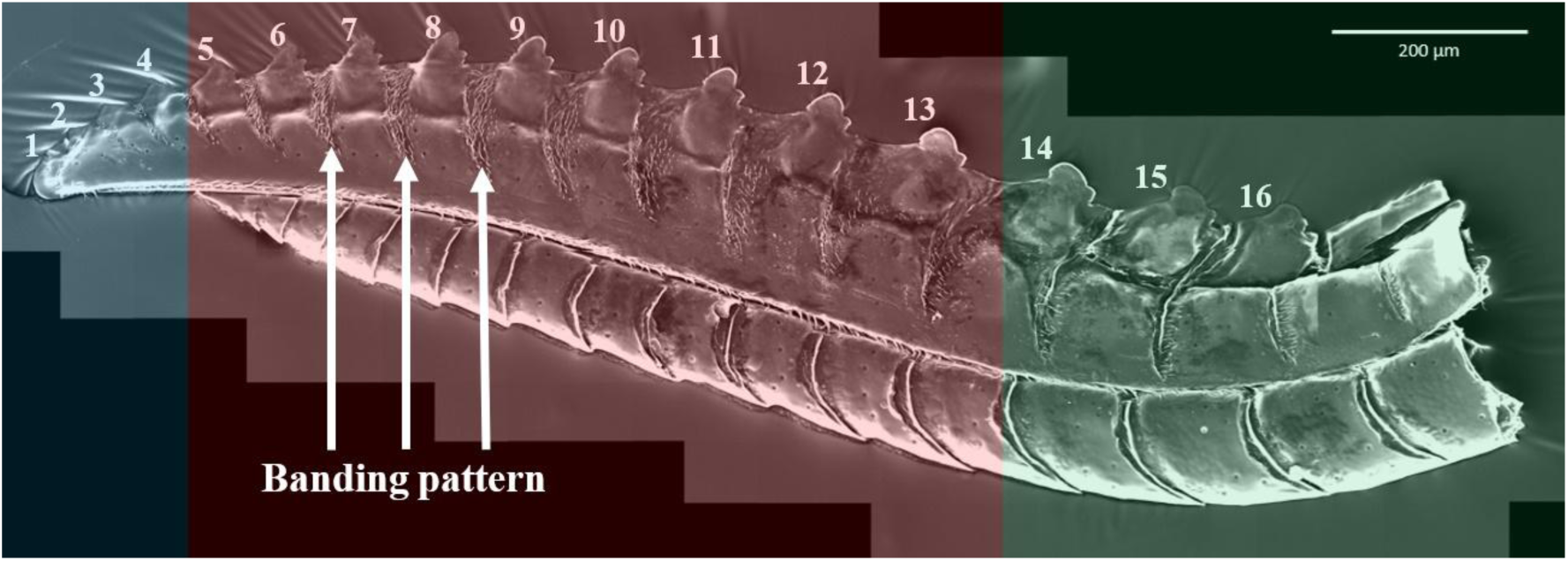
SEM image of R. scalaris ovipositor with tooth numbering and banding pattern. Colours match functional zones. Adapted from [14]. Open access (CC-BY 4.0).

The first four teeth, in the blue area, are involved in the insertion of the ovipositor. These teeth lack both serrulae and bump since the SCM is not required for this phase. The central part of the ovipositor, coloured in red, cuts through cells using the SCM to avoid damaging the vascular bundles. All teeth in this area have a bump. The serrulae are gradually more present as the base of the ovipositor is approached and eventually transform into a bump, thus resulting in teeth with bumps on both sides at the end of the red zone. The angles of the teeth in this area increase gradually from approximately 36° to 66° with proximity to the ovipositor base. This reflects the increasing likelihood of interaction with the outer layer of the plant. In the green zone, the tooth angle jumps from 66° to 90° for these teeth already have bumps on both sides. These teeth engage only with the outmost layer of plant tissue and are not at risk of damaging vascular bundles. The most basal tooth likely does not reach the substrate and may be vestigial. Figure 22 illustrates the correspondence between teeth morphology and plant tissue.

**Figure 22:**
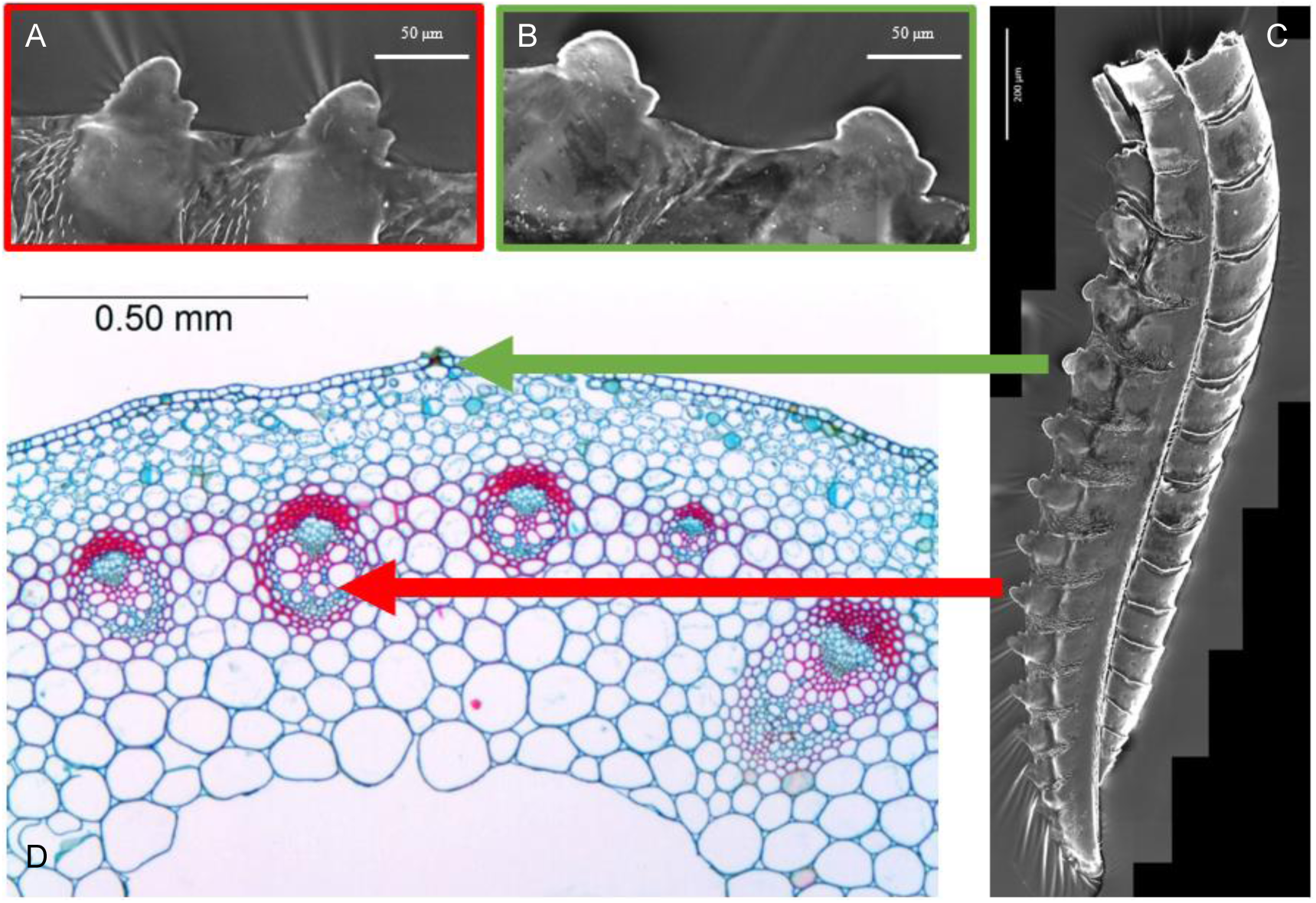
Relation between teeth angle and plant tissue. D: cross-section of Ranunculus acris, adapted with permission from the author [32]. The areas presenting red cells are the vascular bundles. C: SEM image of R. scalaris ovipositor. The green arrow connects the more destructive teeth (B) to the outer layers of the plant. The red arrow connects the teeth capable of selective cutting (A) to the area where vascular bundles are present. Reproduced from [14]. Open access (CC-BY 4.0).

The influence of the tooth angle is particularly relevant in relation to the banding pattern. This banding pattern is a series of stripes running perpendicular to the edge of the blade between adjacent teeth, as shown in Figure 21. In the profilometry of a *R. scalaris* lancet, shown in Figure 15, the bands appear as depressions relative to their surroundings.

Confocal laser scanning microscopy (CLSM) of an *R. scalaris* ovipositor, as seen in Figure 23, shows that the banding pattern has a different material composition from the surrounding rib structure, as they emit different wavelengths when excited.

**Figure 23:**
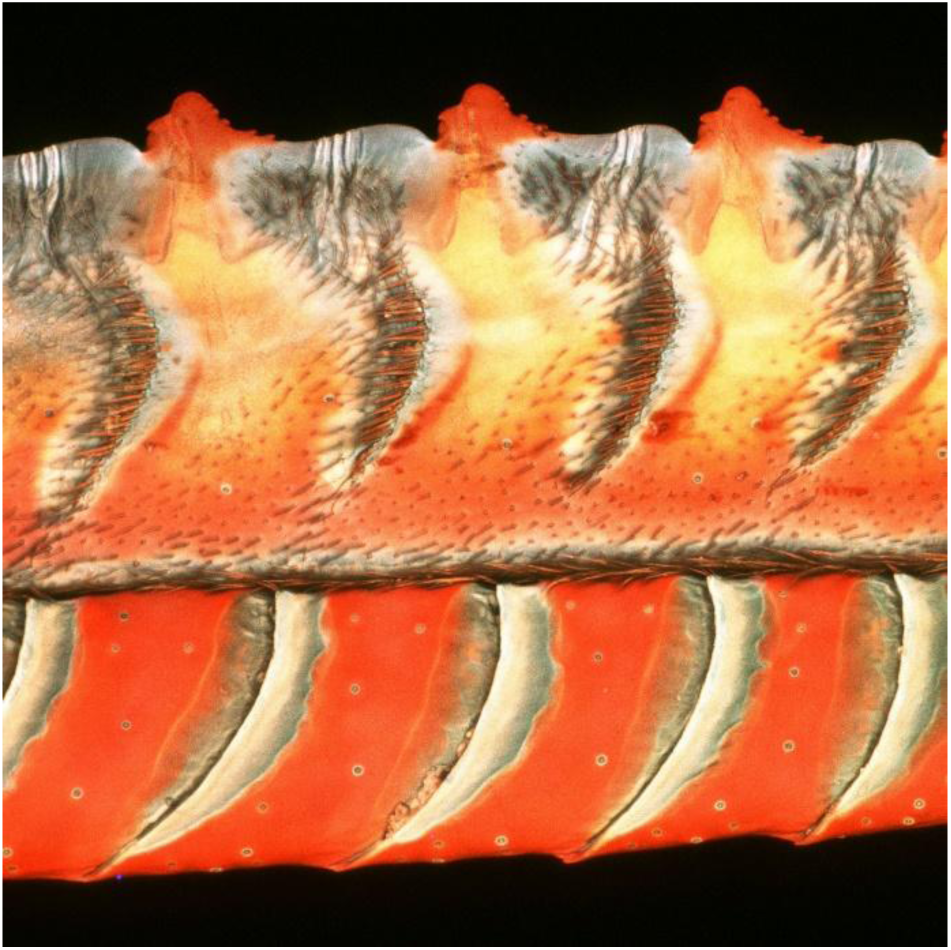
CLSM image of ovipositor of R. scalaris.

Studies on insect cuticle have shown that the wavelength of laser-induced autofluorescence correlates with the elastic modulus of the material [33]. Wavelengths on the blue end of the spectrum indicate lower elastic modulus, while those closer to the red end of the spectrum correspond to higher elastic modulus. A banding pattern consisting of thinner, more compliant material than the teeth may act as a spring-like mechanism connecting adjacent teeth. During reciprocating motion, when a substrate is caught between a tooth from each lancet moving in opposite directions, the resulting stresses affect each blade differently. Figure 24 schematically illustrates how, on the pushing blade (A), the stresses compress the band basal to the pushing tooth, whereas on the pulling blade (B), the band basal to the tooth pulling is stretched. Greater deformation of the band on the pushing blade leads to a smaller tooth angle, resulting in a lower *σth*, meaning that the ejection mechanism is more easily triggered. On the pulling blade, band deformation reduces the attack angle of the bump, lowering the stress on the bump and reducing the likelihood of damaging the substrate. This dynamic reduction in attack angle, shown in Figure 24, is analogous to a reduction in the rake angle on a machining tool.

**Figure 24:**
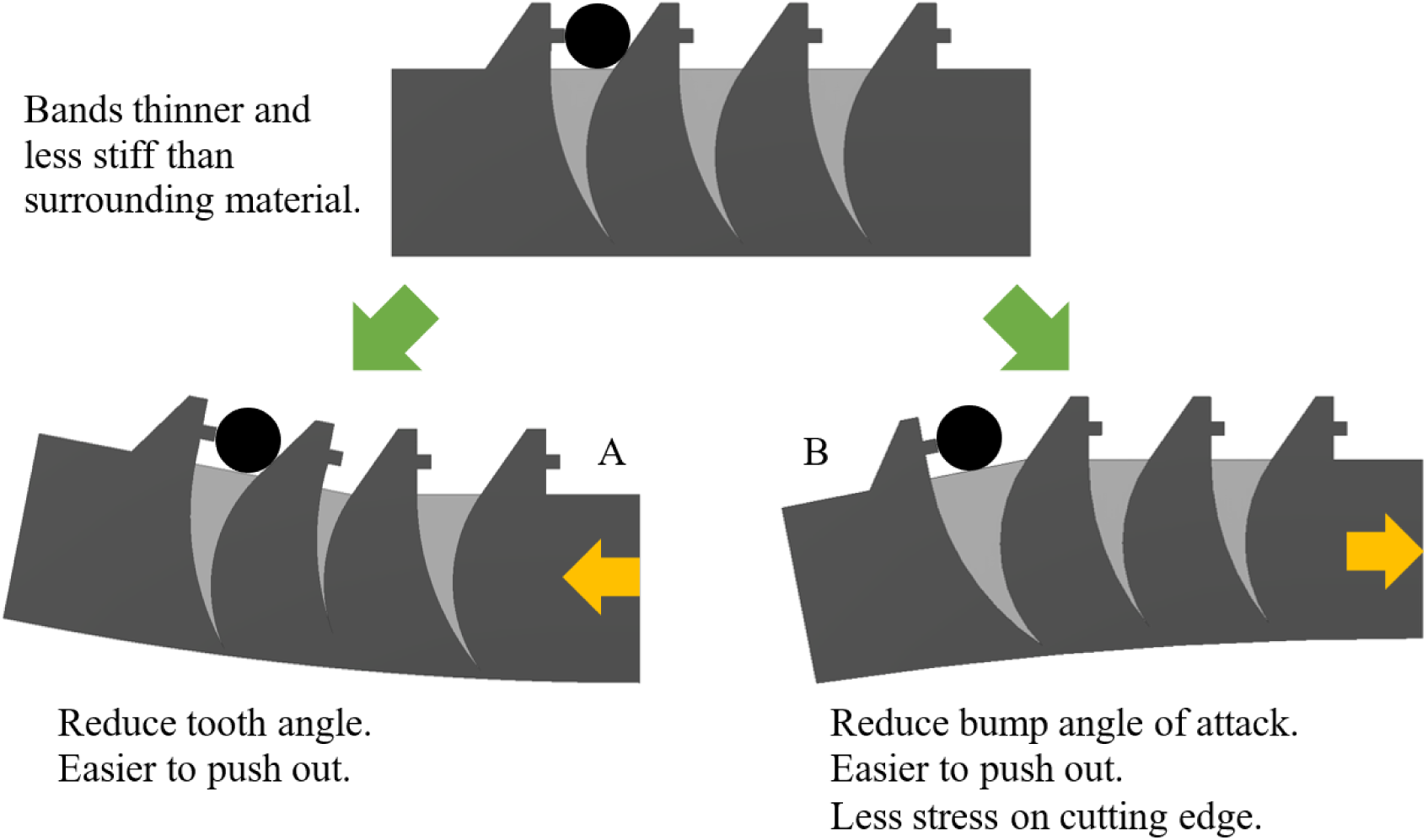
Banding pattern deformation during cutting: compression (A) vs. tension (B).

### 4.3 Complex selective cutting mechanism

Incorporating both modifiers into the SCM increases the complexity of the principle but enhances its effectiveness. The basic SCM discerns substrates based solely on ultimate stress, which is already a significant advantage. The two modifiers add sensitivity to additional material properties: hardness and elastic modulus. Whether the serrulae pierce and hook the substrate depends on its hardness. The banding pattern modifier depends on the elastic modulus of the substrate in relation to that of the bands. During cutting, both the substrate and the bands deform, and their relative stiffness determines which one undergoes larger deformations. Therefore, the complex selective cutting mechanism (CSCM) can discern materials based on ultimate stress, hardness, and elastic modulus, as schematically illustrated in Figure 25.

**Figure 25:**
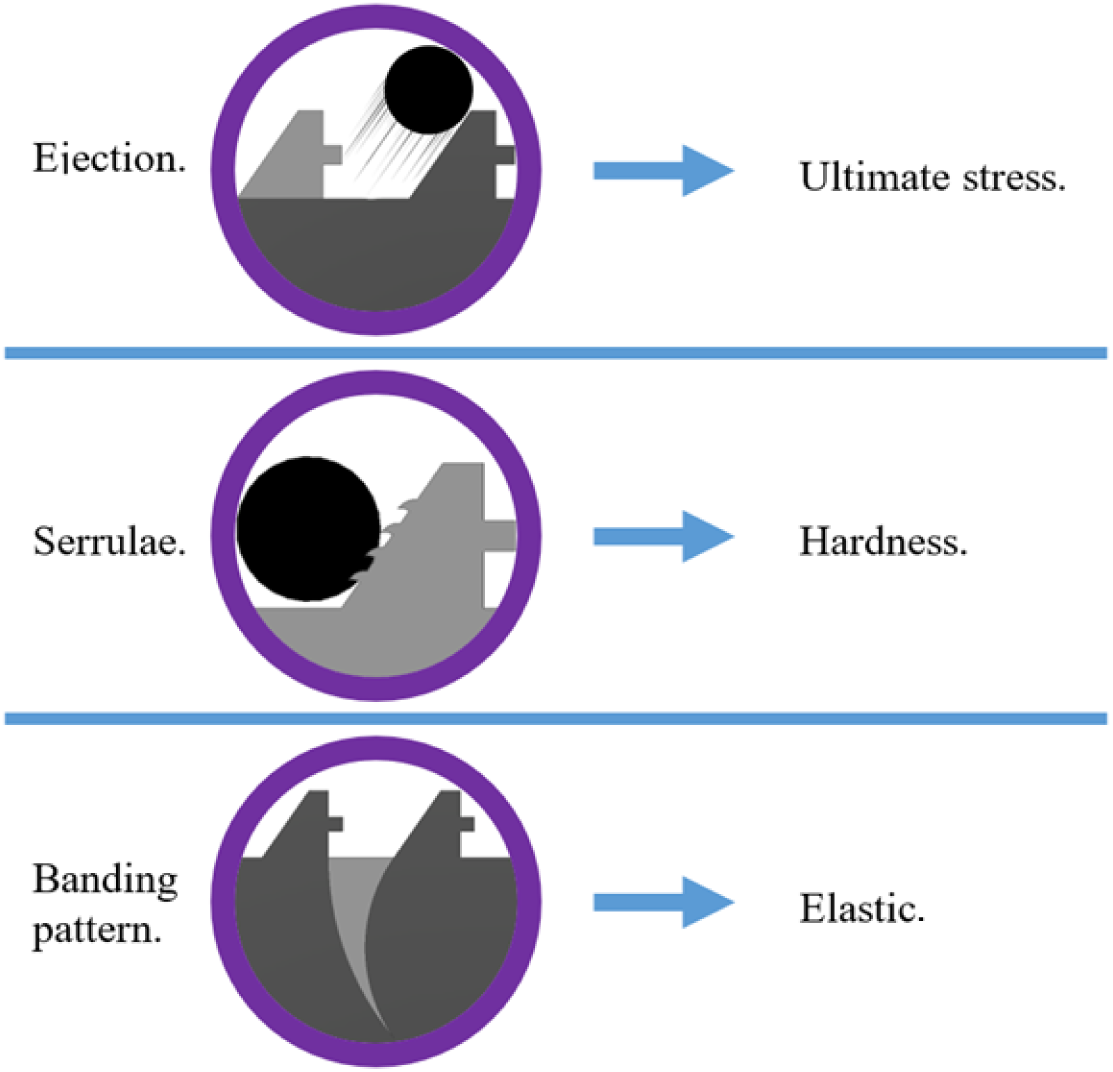
Conceptual summary of the CSCM components and their associated substrate properties.

Adding hardness and stiffness as filtering criteria increases the mechanism’s capacity to distinguish between materials. Even if multiple substrates share the same value for one or two properties, the remaining material properties still allow differentiation. Practically, if tailoring a set of blades to differentiate two materials based on a single property proves difficult, incorporating the other two properties may enhance the specificity of the SCM.

## 5 Limitations and directions for future work

The basic model presented is a first-order approximation to predict *σth* values. While it may already assist in the design of tailored blades, it may not yet be sufficient for applications requiring differentiation between substrates with similar mechanical properties. Its effectiveness could be improved by fully including in the model the two modifiers presented in Section 4. The serrulae effect is included but material hardness is not, as the model does not account for changes in friction coefficient depending on whether the serrulae pierce the substrate. Similarly, the angle effect is represented in the model, but not the banding pattern itself. As a result, the elastic modulus is not a factor unless the user precomputes the band deformation and manually adjusts the angle value.

The model does not account for the speed of the reciprocating motion, since it is considered as a quasi-static system. In many materials, increasing the loading speed results in higher measured ultimate stress. This effect may result in the system being unable to cut materials at the model-predicted *σth*, even when all other variables remain constant. Alternatively, higher loading speed may strengthen the material in regions pierced by the serrulae, preventing local failure and disabling thus the ejection mechanism. Although the speed effect is not considered in the current model, it could be implemented and potentially act as a modifier in the CSCM. A tool capable of regulating the reciprocating motion speed could, in principle, have its *σth* adjusted in real-time. Conversely, a very low loading speed may induce creep in some materials, leading to larger deformations than expected, stress relaxation, and therefore reduced effectiveness in cutting and ejection.

The model does not account also for changes in contact area between the blades and the substrate. Under a constant applied force, the deformation of the substrate may increase the contact area, and therefore reduce the stresses transmitted to the substrate. The resulting lowering of the *σth* due to the stress reduction would be most pronounced in substrates with lower elastic modulus and/or high ductility.

As introduced earlier in the paper, the appropriate selection of a failure criterion is critical to accurately model the behaviour of the system. This can pose a limitation, particularly when data on the material’s mechanical properties are unavailable. Similarly, deciding between the compression and shear versions of the model depends on the available information about the substrates. In the compression model, stress is independent of substrate size, so tooth spacing is only relevant in determining whether the substrate fits between the teeth and can therefore be damaged. If a cellular tissue is considered, larger cells may not fit between the teeth while smaller cells might. In contrast, the shear model calculates lower stress for the same cut force when the substrate is larger. As a result, the substrate size directly influences the *σth* in the shear model. According to the model, larger cells would experience lower stresses than smaller ones. This relevance of the substrate size could be exploited as a filtering modifier in either version of the CSCM.

Individual cells do not behave like a solid block of material. Cells are mechanically an outer membrane or wall filled with liquid. Although for many applications they can be approximated as a material with an ultimate stress, a failure criterion representing them as a fluid-filled membrane could improve the accuracy of the model. For this purpose, the hydrostatic and deviatoric stress tensors already provided by the current model could be useful. Fitting the teeth on each side of the substrate might be seen as challenging, but inner tissue cells are not usually packed in ordered rows, so teeth positioning is unlikely to be problematic. Additionally, since the teeth are much thinner than the cells, friction against the adjacent cells is not an issue. Thus, those two factors are not needed in the model.

## 6 Conclusions

This article explored the SCM based on the ovipositor of sawflies, particularly *Rhogogaster scalaris*. Some sawflies use this mechanism to avoid damaging vital structures while incising plant tissue to lay their eggs. A novel analytical model was developed that captures the core mechanism and reveals additional functional elements linked to morphological traits. The goal is to enable the design of bioinspired tools capable of discriminating substrates based on their mechanical properties, by ejecting those meant to remain undamaged.

The model addresses the SCM as a race to failure or ejection of substrate material depending on which conditions are met first. An equilibrium of forces provides the basis for the ejection criterion, and the stress state enables the application of a failure criterion. The model accommodates multiple failure criteria and two stress formulations: compressive and shear. Based on substrate mechanical properties, applied forces and tooth geometry, the model predicts whether a substrate will be ejected or damaged. This passive selection reproduces the biological principle, which also requires no sensing or feedback.

The model was validated against experimental data from literature where the SCM was tested on manufactured soft substrates [14]. The outcomes of the tests were successfully predicted as a function of force against the substrate, cut force and blade geometry. This includes experiments demonstrating the effect of the serrulae, the bump and the blade scale.

Although the model captures the core principle of the SCM, it also uncovered the influence of additional morphological traits. The serrulae modifier, based on whether the serrulae pierce the substrate or not, can increase the likelihood of substrate failure or ejection respectively. Similarly, an increase in tooth angle reduces the likelihood of substrate ejection. The banding pattern morphological trait is hypothesized to enable dynamic modulation of teeth angles, which in combination with the tooth angle effect results in the banding pattern modifier. While the primary principle of the SCM discriminates substrates based on ultimate strength, the serrulae and banding pattern modifiers introduce sensitivity to hardness and relative elastic modulus, respectively.

The model and the novel selective cutting principles it revealed can inform the development of selective cutting tools based on the mechanical properties of the substrate. This has potential applications in fields such as biomedicine, soft robotics, material engineering or food technologies. The ability to discriminate substrates without sensors or feedback is a robust, yet cognitively undemanding strategy for material differentiation. Despite current limitations, this study provides a transferable insight into how nature achieves a passive CSCM and lays the foundation for the development of future and smarter cutting tools.

## Acknowledgments

One of the authors, Dr Martí Verdaguer Mallorquí, acknowledges the financial support of Heriot-Watt University through the James Watt Scholarship. We thank the Senckenberg Deutsches Entomologisches Institut for providing the sawfly specimens. We thank Mihai Costea for granting permission to adapt the image of *Ranunculus acris*.

## Notes

https://doi.org/10.5281/zenodo.15639471

### Competing Interest Statement

The authors have declared no competing interest.

